# Laterally transferred macrophage mitochondria act as a signaling source promoting cancer cell proliferation

**DOI:** 10.1101/2021.08.10.455713

**Authors:** Chelsea U. Kidwell, Joseph R. Casalini, Soorya Pradeep, Sandra D. Scherer, Daniel Greiner, Jarrod S. Johnson, Gregory S. Olson, Jared Rutter, Alana L. Welm, Thomas A. Zangle, Minna Roh-Johnson

**Author notes:** These authors contributed equally to this work (Chelsea U. Kidwell, Joseph R. Casalini).

## Abstract

Lateral transfer of mitochondria occurs in many physiological and pathological conditions. Given that mitochondria provide essential energy for cellular activities, mitochondrial transfer is currently thought to promote the rescue of damaged cells. We report that mitochondrial transfer occurs between macrophages and breast cancer cells, leading to increased cancer cell proliferation. Unexpectedly, transferred macrophage mitochondria are dysfunctional, lacking mitochondrial membrane potential. Rather than performing essential mitochondrial activities, transferred mitochondria accumulate reactive oxygen species which activates ERK signaling, indicating that transferred mitochondria act as a signaling source that promotes cancer cell proliferation. We also demonstrate that pro-tumorigenic M2-like macrophages exhibit increased mitochondrial transfer to cancer cells. Collectively, our findings reveal how mitochondrial transfer is regulated and leads to sustained functional changes in recipient cells.

**One-Sentence Summary:** Lateral transfer of macrophage mitochondria acts as a ROS signaling source, regulating cancer cell proliferation through ERK signaling.

## Main Text

Recent studies reveal that mitochondrial lateral transfer, the movement of mitochondria from one cell to another, can affect cellular and tissue homeostasis *(1, 2)*. Most of what we know about mitochondrial transfer stems from bulk cell studies and have led to the paradigm that functional transferred mitochondria restore bioenergetics and revitalize cellular functions to recipient cells with damaged or non-functional mitochondrial networks (*3*). However, mitochondrial transfer also occurs spontaneously between cells with functioning endogenous mitochondrial networks and enhances energy-consuming processes such as invasion and proliferation in the recipient cells. The mechanisms underlying how transferred mitochondria can promote such sustained behavioral reprogramming remain unclear.

### Cancer cells with macrophage mitochondria exhibit increased proliferation

We previously reported that macrophages transfer cytoplasmic contents to cancer cells *in vitro* and *in vivo* (*4*), and hypothesized that a macrophage/cancer cell system would be ideal for probing mitochondrial transfer in cells with functioning mitochondrial networks. Our studies employed blood-derived human macrophages and a human breast cancer cell line, MDA-MB-231 (231 cells), stably expressing a mitochondrially localized mEmerald or red fluorescent protein (mito-mEm or mito-RFP, respectively) (**Fig. 1A**). We observed mitochondrial transfer from macrophages to 231 cells using live cell confocal microscopy (**Fig. 1B,** arrowheads) and flow cytometry (**Fig. 1C–D,** flow cytometry scheme in **Fig. S1A**). A range of transfer efficiencies were observed, which we attribute to donor-to-donor variability (Fig. 1D). We also observed mitochondrial transfer to a non-malignant breast epithelial cell line, MCF10A, albeit at lower rates (**Fig. S1B**). Transferred mitochondria contain a key outer mitochondrial membrane protein, TOMM20 (**Fig. S1C**, arrowhead) and mitochondrial DNA (**Fig. S1D**, arrowhead), suggesting that intact organelles are transferred to 231 cells. To define the requirements for transfer, we performed trans-well experiments in which we cultured 231 cells either physically separated from macrophages by a 0.4μM trans-well insert or in contact with macrophages (scheme in **Fig. S1E**), or with conditioned media (**Fig. S1G**). These data showed that mitochondrial transfer increased dramatically under conditions where 231 cells could contact macrophages directly (**Fig. S1F-G**). Due to the low rates of mitochondrial transfer across macrophage donors (0.84%, Fig. 1D), we subsequently took advantage of single cell, high-resolution approaches – rather than bulk approaches – to follow the fate and functional status of transferred mitochondria.

**Fig. 1.**
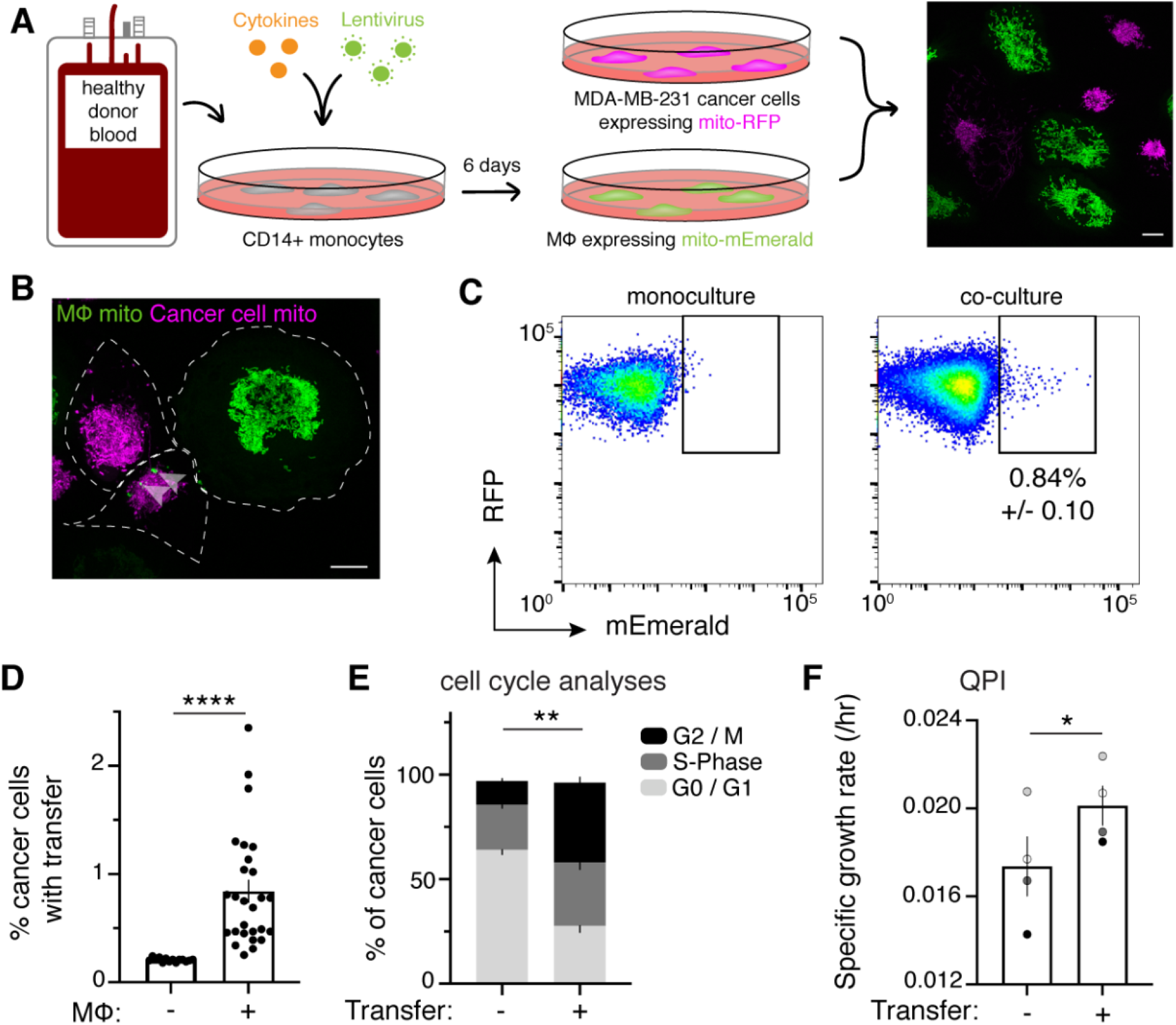
Cell contact-mediated transfer of macrophage mitochondria leads to increased cancer cell proliferation. (A) CD14+ monocytes harvested from human blood are transduced and differentiated for 6 days. Mito-mEm+ macrophages (green) are co-cultured with MDA-MB-231 cells (231 cells) expressing mito-red fluorescent protein (mito-RFP; magenta; right image). (B) Confocal image showing transferred mitochondria (green, arrowhead) in a 231 cell (magenta, cell outline in white). (C) Representative flow cytometry plots depicting mitochondrial transfer (black box) within a population of co-cultured mito-RFP+ 231 cells (right) compared to monoculture control (left) with background level of mEmerald fluorescence set at 0.2%. (D) Aggregate data of mitochondrial transfer rates across macrophage donors. Each data point represents one replicate (N=14 donors). (E) Ki-67 levels and DNA content was measured in co-cultured 231 cells after 24 hours. Percentage of cancer cells within a specific cell cycle phase with or without transfer is shown. A significantly different percent of recipient cells occupies G2 and M (black) compared to non-recipient cells (N=5 donors; statistics for G2/M only). (F) Co-cultured recipient 231 cells have a significantly higher specific growth rate compared to non-recipients (N=4 donors indicated as shades of gray). For all panels, standard error of the mean (SEM) is displayed and scale bars are 10 μm. Unpaired t-test with Welch’s correction, *p<0.05; **p<0.01; ***p<0.0001.

To determine the effects of macrophage mitochondrial transfer on cancer cells, we performed single cell RNA-sequencing on cancer cells that received macrophage mitochondria. These data revealed that mitochondrial transfer induced significant changes in canonical cell proliferation-related pathways (**Fig. S2A**). To follow up on the RNA-sequencing results, we used flow cytometry to evaluate cell cycle changes, and found significant increases in the percent of cells within the G2 and Mitotic (M) phases of the cell cycle in recipient cells, as compared to their co-cultured counterparts that did not receive mitochondria (**Fig. 1E,** flow cytometry scheme in **Fig. S2B, Fig. S2C-D**). These cells were not undergoing cell cycle arrest, as we found that recipient cells completed cytokinesis at rates equivalent to their co-cultured non-recipient counterparts (**Fig. S2E**). For further confirmation of this proliferative phenotype, we used quantitative phase imaging (QPI) to detect changes in dry mass of co-cultured 231 cells over time (*5*). Consistent with the flow cytometry-based cell cycle analysis, the specific growth rates increased significantly in 231 cells with macrophage mitochondria compared to 231 cells that did not receive mitochondria (**Fig. 1F**). Using QPI, we also measured the growth rates of daughter cells born from recipient 231 cells containing macrophage mitochondria (*6*). Daughter cells that inherited the “mother’s” macrophage mitochondria exhibited an increase in their rate of change of dry mass over time versus sister cells that did not inherit macrophage mitochondria (**Fig. S3A-C**). These data indicate the enhanced proliferation observed after mitochondrial transfer is not simply due to proliferative cells more readily receiving transfer. Furthermore, these results indicate that mitochondrial transfer promotes a sustained pro-growth and proliferative effect in both recipient and subsequent daughter cells.

### Transferred mitochondria accumulate reactive oxygen species in cancer cells

We next sought to understand how donated mitochondria can stimulate a proliferative response in recipient cells. We performed time-lapse confocal microscopy on co-cultures and found that macrophage-derived mito-mEm+ mitochondria remained distinct from the recipient host mitochondrial network, with no detectable loss of the fluorescent signal for over 15 hours (**Fig. 2A**, arrowhead; **Movie S1**). To probe the functional state of the donated mitochondria, we performed live imaging with MitoTracker Deep Red (MTDR), a cell-permeable dye that is actively taken up by mitochondria with a membrane potential (*7*). To our surprise, all of the transferred mitochondria were MTDR-negative (**Fig. 2B**, top left). This was also confirmed using a different mitochondrial membrane potential-sensitive dye, Tetramethylrhodamine Methyl Ester (TMRM; Fig. S1D). We also labeled lysosomes and acidic vesicles with a dye, LysoTracker, and found that the majority of transferred mitochondria (57%) did not co-localize with the LysoTracker signal (Fig. 2B, top right). The status of transferred mitochondria was unexpected because mitochondria typically maintain strong membrane potentials, and dysfunctional mitochondria that lack membrane potential are normally degraded or repaired by fusion with healthy mitochondrial networks (*8*). In addition, we confirmed that 91% of transferred mitochondria were not encapsulated by a membranous structure, thus excluding sequestration as a mechanism for explaining the lack of degradation or interaction with the endogenous mitochondrial network (**Fig. S4A**).

**Fig. 2.**
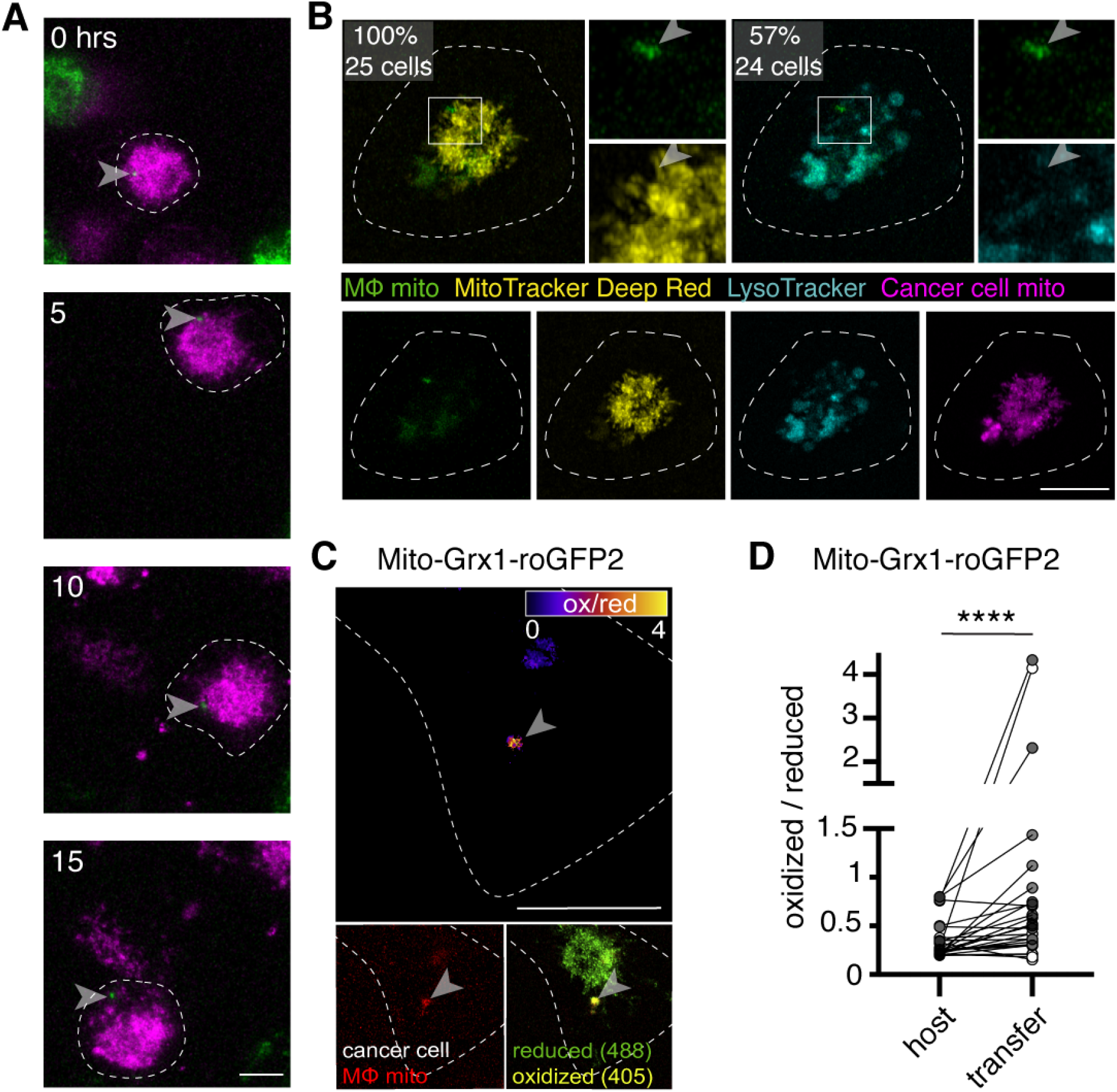
Transferred macrophage mitochondria are long-lived, depolarized, and accumulate reactive oxygen species. (A) Stills from time-lapse imaging depicting the longevity of the transferred mitochondria (green, arrowhead) within a 231 cell (magenta, cell outline in white). Time elapsed listed in left corner. (B) Confocal image of a mito-RFP+ 231 cell (magenta) containing macrophage mitochondria (green, arrowhead) stained with MTDR (yellow) and LysoTracker (teal). MTDR does not accumulate in 100% of donated mitochondria (N=25 cells, 5 donors). Majority (57%) of donated mitochondria do not colocalize with LysoTracker signal (N=24 cells, 4 donors). (C) Ratiometric quantification of mito-Grx1-roGFP2 biosensor mapped onto the recipient 231 cell with fire LUT (top panel). Confocal image of mito-Grx1-roGFP2-expressing 231 cell (bottom right, green and yellow) containing a macrophage mitochondria (bottom left, red, arrowhead). (D) Ratiometric measurements of the mito-Grx1-roGFP2 sensor per 231 cell (paired dots) at a region of interest containing the host mitochondrial network (host) or a transferred mitochondria (transfer). Cells were co-cultured for 24 hours (N=27 cells, 3 donors indicated in shades of gray). Scale bars are 10 μm. Wilcoxon matched-pairs signed rank test, ****p<0.0001.

Our work indicates that transferred macrophage mitochondria are depolarized but remain in the recipient cancer cell. Therefore, we hypothesized that donated mitochondria act as signal source rather than energy. One signal associated with mitochondria is reactive oxygen species (ROS), which normally occur as byproducts of mitochondrial respiration, and can be produced at high levels during organellar dysfunction (*9*). Using a genetically encoded biosensor, mito-Grx1-roGFP2, as a live readout of the mitochondrial glutathione redox state (*10*), we found that after 24 and 48 hours, significantly higher ratios of oxidized:reduced protein were associated with the transferred mitochondria versus the host network (**Fig. 2C–D**, **S4B**). These data indicate that transferred macrophage mitochondria in recipient cells are associated with higher levels of oxidized glutathione, suggesting that they are producing higher amounts of ROS. Consistent with these results, a second biosensor that is specific for the ROS H_2_O_2_, mito-roGFP2-Orp1 (*11*), also reported more oxidation at the transferred mitochondria compared to the host network (**Fig. S4C-D**) after 48 hours of co-culture. At 24 hours, we observed a similar trend, but no statistically significant difference (Fig. S4D). These results indicate ROS accumulates at the site of transferred mitochondria in recipient cancer cells.

### ROS accumulation leads to ERK-dependent recipient cell proliferation

To test whether ROS accumulation can induce cancer cell proliferation, we stably expressed a mitochondrially localized photosensitizer, mito-KillerRed, which generates ROS when photobleached with 547nm light (*12*). As expected, photobleaching mito-KillerRed+ regions of interest induced ROS (*13*) (**Fig. S5A**) and also significantly increased the rate of cell division compared to control (mito vs. cyto bleach; **Fig. 3A, S5B-C**). These results indicate that ROS induction can promote cancer cell proliferation. ROS induces several downstream signaling pathways (*9, 14*). Consistent with this regulation, we found a significant increase in ERK signaling in recipient cancer cells, as assayed using the ERK-Kinase Translocation Reporter (ERK-KTR) (*15*), which translocates from the nucleus to the cytoplasm when ERK is activated (ERK-KTR quantification described in **Fig. S6-7**). Recipient 231 cells had significantly higher cytoplasmic to nuclear (C/N) ERK-KTR ratios compared to cells that did not receive transfer, indicating increased ERK activity (**Fig. 3B–C, S8A-B**).

**Fig. 3.**
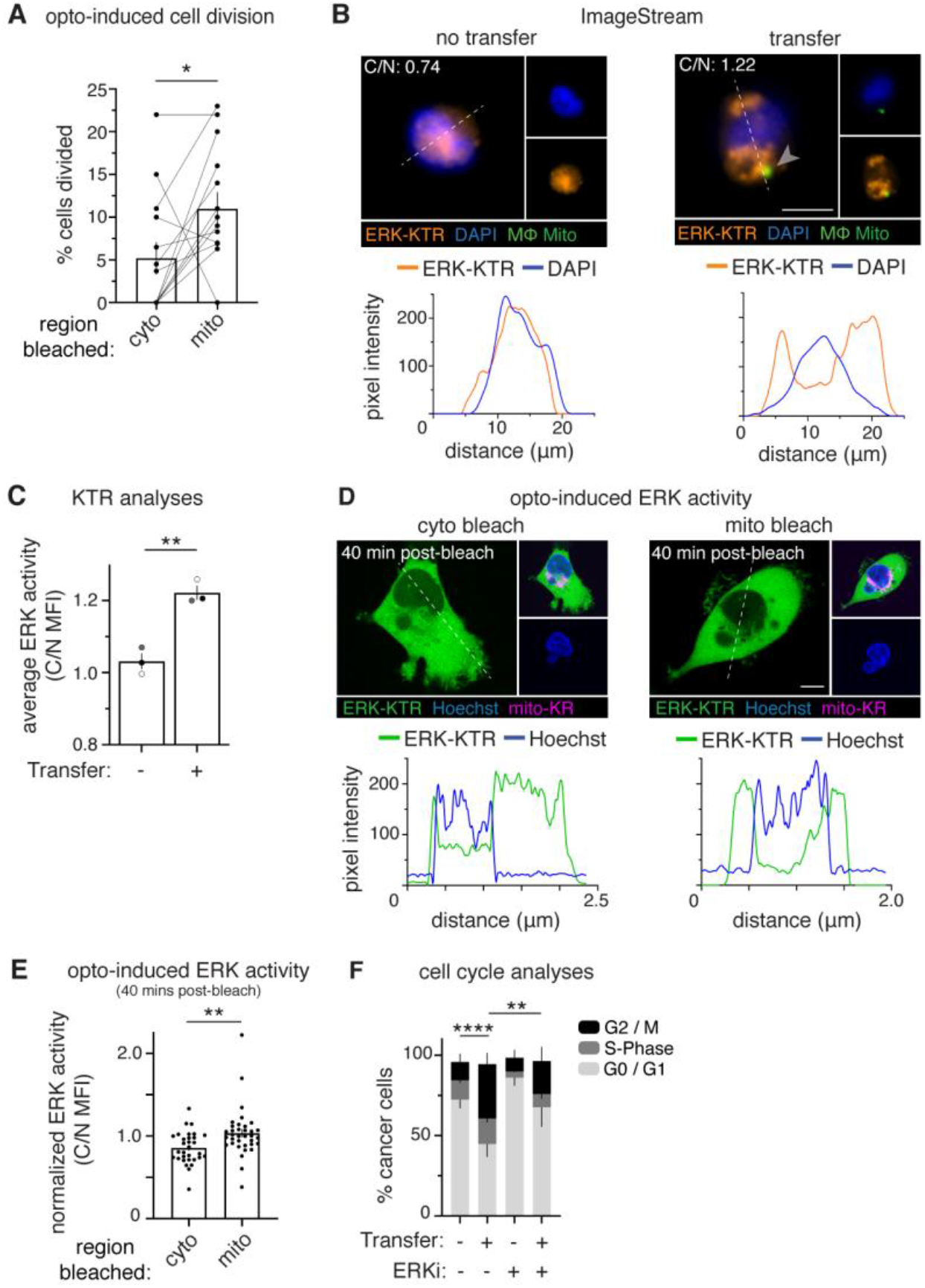
Recipient cancer cells exhibit ERK-dependent proliferation. (A) Quantification of cell division after photobleaching. Each data point is the average within a field of view (N=13 experiments), with control (cyto) and experimental (mito) data shown as paired dots per experiment. (B) ImageStream was used to measure the MFI of an ERK-Kinase Translocation Reporter (ERK-KTR, orange) in the nucleus (DAPI, blue) or cytoplasm of co-cultured 231 cells that did (right) or did not (left) receive mitochondria (green, arrowhead). Below: representative line scans (white dotted lines) of ERK-KTR (orange) and DAPI (blue). (C) Average ERK activity from data displayed in (D) (cytoplasm/nucleus (C/N) mean fluorescence intensity (MFI); N=3 donors indicated as shades of gray). (D) Confocal images of 231 cells expressing ERK-KTR (green) and Mito-KillerRed (magenta) with Hoechst 33342 (blue), after control cytoplasmic bleach (cyto, left) or mito-KillerRed+ bleach (mito, right). Below: representative line scans (white dotted lines) of ERK-KTR (green) and DAPI (blue). (E) Quantification of ERK-KTR translocation 40 minutes post-bleach (cyto vs. mito), normalized to time 0. (F) Cell cycle analyses of co-cultured 231 cells treated with vehicle or ERK inhibitor (ERKi) with or without transfer (N=3 donors; statistics for G2/M only). Error bars represent SEM and scale bars are 10 μm. Wilcoxon matched-pairs signed rank test (A), Unpaired t-test (C, E), 2-way ANOVA (F), *p<0.05; **p<0.01; ****p<0.0001.

We next asked whether recipient cell proliferation was caused by ROS-induced ERK activation. By expressing both the mito-KillerRed and ERK-KTR in 231 cells, we show that photobleaching KillerRed+ mitochondria induced ERK-KTR translocation, indicating that ROS induction is sufficient to increase ERK activity in cancer cells (**Fig. 3D–E, S8C**). Furthermore, treatment with an ERK inhibitor (ERKi) was sufficient to inhibit ERK activity (**Fig. S9A-B**) and specifically decreased proliferation of recipient 231 cells versus control (**Fig. 3F, S9C**), providing further evidence that mitochondrial transfer increases ERK-dependent proliferation. As expected, ERKi treatment did not alter mitochondrial transfer efficiencies, showing that ERK signaling does not influence mitochondrial transfer (**Fig. S9D**). Together, these results indicate that cancer cells that receive macrophage mitochondria proliferate in response to ROS accumulation in an ERK-dependent manner.

### Pro-tumorigenic M2-like macrophages transfer more mitochondria to patient-derived cells

In many solid tumors, macrophages alter their differentiation status depending on environmental stimuli (*16*). Therefore, we next tested how macrophage differentiation status affects mitochondrial transfer using pro-inflammatory M1-like and pro-tumorigenic M2-like macrophage subtypes (*17*) (**Fig. S10A**). We observed that mitochondria that were transferred from both M1-like and M2-like macrophages were depolarized in recipient 231 cells (**Fig.S10B**). However, M2-like macrophages exhibited increased mitochondrial fragmentation (**Fig. 4A–B**, **S10C-E**) and transferred mitochondria to 231 cells more efficiently (**Fig. 4C**) than did M1-like and M0 (unstimulated) macrophages. Given this observation, we hypothesized that smaller mitochondrial fragments might be transferred more readily than larger networks. To test this hypothesis, we directly manipulated mitochondrial morphology by modulating a key regulator of mitochondrial fission, DRP1 (*18*). Macrophages which expressed either *DRP1*-shRNA (hyper-fused networks) or mCherry-DRP1 overexpression (hyper-fragmented networks) (**Fig. 4D**, **S10F-G**) showed decreased or increased mitochondrial transfer, respectively (**Fig. 4E**), indicating that mitochondrial morphology can alter the efficiency of mitochondrial transfer. These data provide a fascinating example of how the intracellular dynamics of one cell can influence how it affects the behavior of a neighboring cell.

**Fig. 4.**
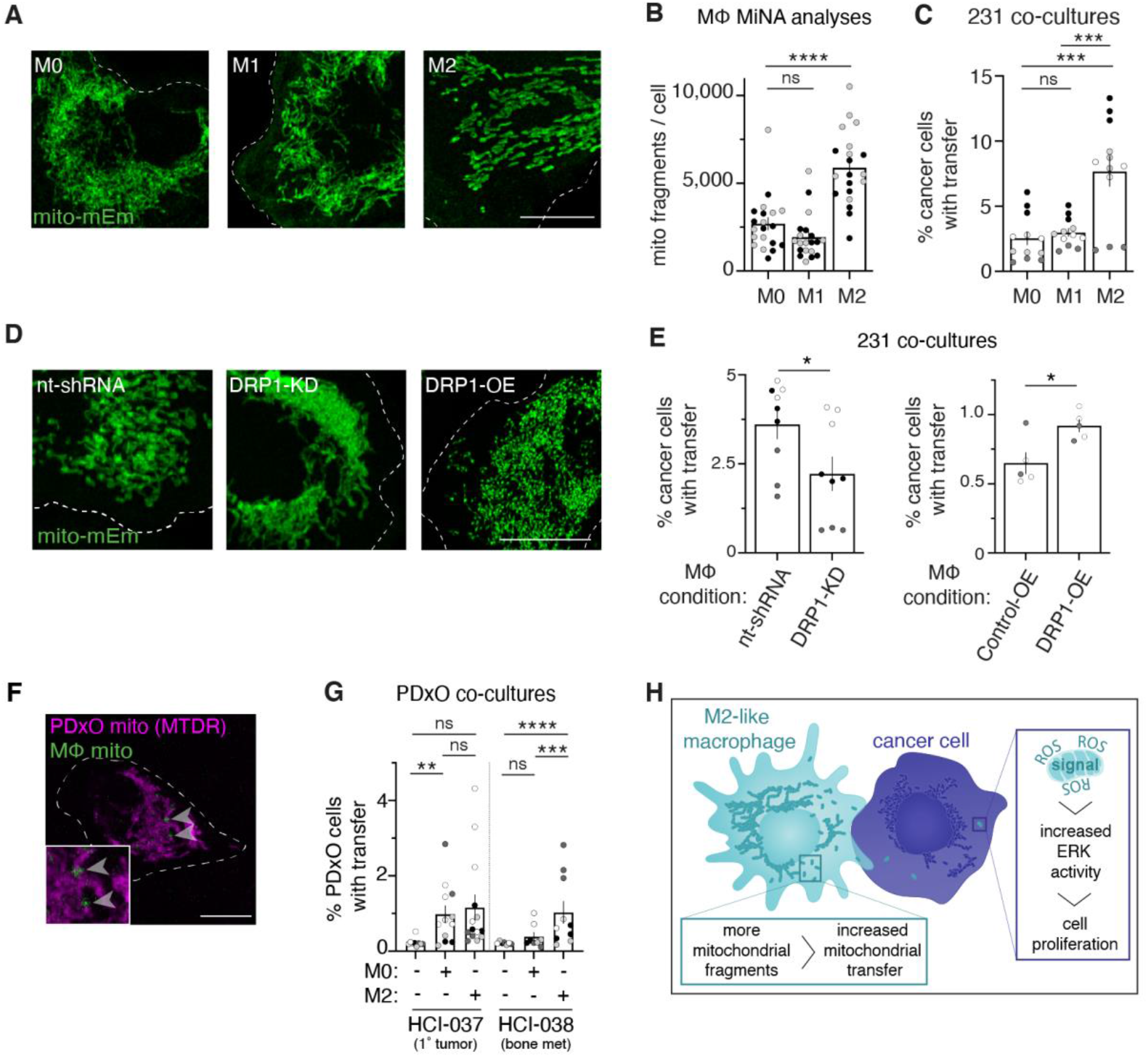
M2-like macrophage activation alters mitochondrial morphology and promotes transfer to breast cancer cells and patient-derived xenograft organoid (PDxO) cells. (A) Representative images of mito-mEm+ macrophages that were non-stimulated (M0, left) or activated to become M1-like (middle) or M2-like (right). (B) Mitochondrial network analyses (MiNA) were used to determine number of mitochondrial fragments per cell (N=2 donors). (C) Macrophages were co-cultured with mito-RFP 231 cells for 24 hours and transfer was quantified with flow cytometry (N=4 donors). (D) Representative images of mito-mEm (green) macrophages with genetic modifications as indicated in upper left corner of each panel. (E) Rates of mitochondrial transfer with DRP1-knockdown (KD; left) or DRP1-overexpression (OE; right) macrophages compared to appropriate controls (N=2 donors). (F) Representative image of a FACS-isolated PDxO cell containing macrophage mitochondria (green, arrowhead) that are MTDR-negative. (G) Rate of mitochondrial transfer to HCI-037 (left 3 columns) or HCI-038 (right 3 columns) PDxO cells from M0 or M2 macrophages (each dot is one replicate, N= 4 donors). (H) Working model for macrophage mitochondrial transfer to breast cancer cells. For all panels, individual donors are indicated as shades of gray with each cell as a data point (except Fig. 4G), SEM is displayed and scale bars are 10 μm. 2-way ANOVA (B-C, G), Unpaired t-test (E), *p<0.05, **p<0.01; ***p<0.001; ****p<0.0001.

To assess whether mitochondrial transfer also occurs in a clinically relevant cancer model, we used three-dimensional stable organoid cultures generated from patient-derived xenografts (PDxOs) (*19*). We examined organoids from a recurrent primary breast tumor (HCI-037) and a bone metastasis (HCI-038) derived from the same breast cancer patient. PDxOs (**Fig. S11A**, top) were dissociated, combined with mito-mEm+ macrophages (Fig. S11A, bottom), and then embedded in Matrigel (experimental scheme in **Fig. S11B**). After 72 hours, mitochondrial transfer was assayed by live imaging (**Fig. 4F)** and quantified with flow cytometry (**Fig. 4G**, flow cytometry scheme in **Fig. S11C**). Mitochondrial transfer was observed from macrophages to both HCI-037 and HCI-038 PDxO cells (Fig. 4G), although intriguingly, M2-like macrophages preferentially transferred mitochondria to the bone metastasis PDxO cells (HCI-038) when compared to primary breast tumor PDxO cells (HCI-037). In all cases, transferred macrophage mitochondria lacked membrane potential (Fig. 4F), consistent with our results in 231 recipient cells. Taken together, our work supports a model (**Fig. 4H**) whereby M2-like macrophages exhibit mitochondrial fragmentation, allowing for enhanced mitochondrial transfer to cancer cells. In the recipient cancer cell, transferred mitochondria are long-lived, depolarized, and accumulate ROS, leading to increased ERK activity and subsequent cancer cell proliferation.

Lateral mitochondrial transfer is a relatively young and rapidly evolving field. Our observation that transferred mitochondria lack membrane potential raises several questions, including when and how transferred mitochondria become depolarized and why they are not repaired or degraded in the recipient cell, given that 231 cells are capable of performing mitophagy (*20*). Impaired mitophagy and enhanced mitochondrial dysfunction are hallmarks of age (*21*), yet little is known about how donor cell mitochondrial health influences mitochondrial transfer. Given that most cancers are age-related diseases, our findings prompt further investigation into how age-associated macrophage mitochondrial dysfunction may contribute to mitochondrial transfer and tumor progression.

Our findings are consistent with several studies describing a metastatic advantage in cancer cells that receive exogenous mitochondria (*22, 23*). However, the mechanism underlying this behavior is unexpected. Studies examining mitochondrial transfer have typically used recipient cells with damaged or non-functional mitochondria, and the fate and function of donated mitochondria are rarely followed in recipient cells. Furthermore, it was largely unclear how transferred mitochondria can affect the behavior of recipient cells with functioning endogenous mitochondrial networks, particularly if the donated mitochondria only account for a small fraction of the total mitochondrial network in the recipient cell. Our work detailing how transferred mitochondria can function as a signaling source provides an explanation for how transferred mitochondria can generate in a sustained behavioral response in recipient cells.

## Acknowledgments

We would like to thank Wes Sundquist and all members of the Roh-Johnson lab for helpful discussions and edits to this manuscript; ARUP Laboratories for providing leukofilters; James Carrington and Joshua Mont for technical support; Hannah Young for help with data analysis; Alan Aderem and Elizabeth Gold for their mentorship to GSO; and Kenneth M. Boucher for help with statistics. We also thank the Huntsman Cancer Institute Cancer Center Shared Resources; the University of Utah Flow Cytometry Core for technical assistance; and the University of Utah Cell Imaging Core for use of the Leica Yokogawa CSU-W1 spinning disc confocal microscope.

## Funding

National Cancer Institute of the National Institutes of Health R37CA247994 (MRJ)
Department of Defense W81XWH-20-1-0591 (MRJ)
The Mary Kay Foundation 10-19 (MRJ)
National Cancer Institute of the National Institutes of Health R00CA190836 (MRJ, CK)
National Cancer Institute of the National Institutes of Health F31CA250317 (JRC)
Department of Defense W81XWH1910065 (TAZ)
National Cancer Institute of the National Institutes of Health U54CA224076 (ALW)
Breast Cancer Research Foundation Founders Fund (ALW)
National Center for Research Resources of the National Institutes of Health 1S10OD026959-01A1 (University of Utah Flow Cytometry Core)
National Cancer Institute of the National Institutes of Health P30CA042014 (Huntsman Cancer Institute Cancer Shared Resources)

## Author contributions

Conceptualization: MRJ, CK, JC
Data Curation: MRJ, CK, JC, SP, DG
Formal analysis: MRJ, CK, JC, SP, TAZ, DG
Funding acquisition: MRJ, ALW, TAZ, CK, JC
Investigation: MRJ, CK, JC, SP, TAZ, DG
Methodology: MRJ, CK, JC, SP, TAZ, JSJ, SDS, ALW
Project administration: MRJ, CK, JC
Resources: MRJ, CK, JC, SP, TAZ, DG, GSO, JSJ, SDS, ALW, JR
Software: SP, TAZ, DG Supervision: MRJ, CK, JC
Validation: MRJ, CK, JC, SP Visualization: MRJ, CK, JC
Writing – original draft: MRJ, CK, JC
Writing – review & editing: MRJ, CK, JC, SP, TAZ, DG, GSO, JSJ, SDS, ALW, JR

## Competing interests

University of Utah may license PDxO models to for-profit companies, which may result in tangible property royalties to members of the Welm lab (SDS and ALW). The other authors declare that they have no competing interests.

## Data and materials availability

The code for QPI analysis is available on GitHub (https://github.com/Zangle-Lab/Macrophage_tumor_mito_transfer).

Single-cell RNA-sequencing data are available in GEO accession number GSE181410. The code for single-cell RNA-sequencing analysis is available on GitHub (https://github.com/rohjohnson-lab/kidwell_casalini_2021).

All other data are available in the main text or the supplementary materials.

## Supplementary Materials

Materials and Methods
Figs. S1 to S11
References (24-33)
Movie S1

## Notes

https://github.com/Zangle-Lab/Macrophage_tumor_mito_transfer

https://www.ncbi.nlm.nih.gov/geo/query/acc.cgi?acc=GSE181410

https://github.com/rohjohnson-lab/kidwell_casalini_2021

## References

1. A. Rustom, R. Saffrich, I. Markovic, P. Walther, H. H. Gerdes, Nanotubular highways for intercellular organelle transport. Science 303, 1007–1010 (2004).

2. D. Torralba, F. Baixauli, F. Sanchez-Madrid, Mitochondria Know No Boundaries: Mechanisms and Functions of Intercellular Mitochondrial Transfer. Front Cell Dev Biol 4, 107 (2016).

3. D. Liu et al., Intercellular mitochondrial transfer as a means of tissue revitalization. Signal Transduct Target Ther 6, 65 (2021).

4. M. Roh-Johnson et al., Macrophage-Dependent Cytoplasmic Transfer during Melanoma Invasion In Vivo. Dev Cell 43, 549–562 e546 (2017).

5. T. A. Zangle, M. A. Teitell, Live-cell mass profiling: an emerging approach in quantitative biophysics. Nat Methods 11, 1221–1228 (2014).

6. T. A. Zangle, M. A. Teitell, J. Reed, Live cell interferometry quantifies dynamics of biomass partitioning during cytokinesis. PLoS One 9, e115726 (2014).

7. M. Poot et al., Analysis of mitochondrial morphology and function with novel fixable fluorescent stains. J Histochem Cytochem 44, 1363–1372 (1996).

8. D. G. Phinney et al., Mesenchymal stem cells use extracellular vesicles to outsource mitophagy and shuttle microRNAs. Nat Commun 6, 8472 (2015).

9. M. Schieber, N. S. Chandel, ROS function in redox signaling and oxidative stress. Curr Biol 24, R453–462 (2014).

10. M. Gutscher et al., Real-time imaging of the intracellular glutathione redox potential. Nat Methods 5, 553–559 (2008).

11. M. Gutscher et al., Proximity-based protein thiol oxidation by H2O2-scavenging peroxidases. J Biol Chem 284, 31532–31540 (2009).

12. M. E. Bulina et al., A genetically encoded photosensitizer. Nat Biotechnol 24, 95–99 (2006).

13. D. A. Bass et al., Flow cytometric studies of oxidative product formation by neutrophils: a graded response to membrane stimulation. J Immunol 130, 1910–1917 (1983).

14. V. Brillo, L. Chieregato, L. Leanza, S. Muccioli, R. Costa, Mitochondrial Dynamics, ROS, and Cell Signaling: A Blended Overview. Life (Basel) 11, (2021).

15. S. Regot, J. J. Hughey, B. T. Bajar, S. Carrasco, M. W. Covert, High-sensitivity measurements of multiple kinase activities in live single cells. Cell 157, 1724–1734 (2014).

16. Y. Pan, Y. Yu, X. Wang, T. Zhang, Tumor-Associated Macrophages in Tumor Immunity. Front Immunol 11, 583084 (2020).

17. X. Huang, Y. Li, M. Fu, H. B. Xin, Polarizing Macrophages In Vitro. Methods Mol Biol 1784, 119–126 (2018).

18. T. B. Fonseca, A. Sanchez-Guerrero, I. Milosevic, N. Raimundo, Mitochondrial fission requires DRP1 but not dynamins. Nature 570, E34–E42 (2019).

19. K. P. Guillen et al., A breast cancer patient-derived xenograft and organoid platform for drug discovery and precision oncology. bioRxiv, 2021.2002.2028.433268 (2021).

20. T. G. Biel, V. A. Rao, Mitochondrial dysfunction activates lysosomal-dependent mitophagy selectively in cancer cells. Oncotarget 9, 995–1011 (2018).

21. G. Chen, G. Kroemer, O. Kepp, Mitophagy: An Emerging Role in Aging and Age-Associated Diseases. Front Cell Dev Biol 8, 200 (2020).

22. L. X. Zampieri, C. Silva-Almeida, J. D. Rondeau, P. Sonveaux, Mitochondrial Transfer in Cancer: A Comprehensive Review. Int J Mol Sci 22, (2021).

23. M. van der Merwe, G. van Niekerk, C. Fourie, M. du Plessis, A. M. Engelbrecht, The impact of mitochondria on cancer treatment resistance. Cell Oncol (Dordr), (2021).

24. J. S. Johnson et al., A Comprehensive Map of the Monocyte-Derived Dendritic Cell Transcriptional Network Engaged upon Innate Sensing of HIV. Cell Rep 30, 914–931 e919 (2020).

25. Y. Hao et al., Integrated analysis of multimodal single-cell data. bioRxiv, 2020.2010.2012.335331 (2020).

26. C. Hafemeister, R. Satija, Normalization and variance stabilization of single-cell RNA-seq data using regularized negative binomial regression. Genome Biol 20, 296 (2019).

27. A. Kramer, J. Green, J. Pollard, Jr., S. Tugendreich, Causal analysis approaches in Ingenuity Pathway Analysis. Bioinformatics 30, 523–530 (2014).

28. J. C. Crocker, D. G. Grier, Methods of digital video microscopy for colloidal studies. J Colloid Interf Sci 179, 298–310 (1996).

29. B. Morgan, M. C. Sobotta, T. P. Dick, Measuring E(GSH) and H2O2 with roGFP2-based redox probes. Free Radic Biol Med 51, 1943–1951 (2011).

30. W. W. Chen, E. Freinkman, T. Wang, K. Birsoy, D. M. Sabatini, Absolute Quantification of Matrix Metabolites Reveals the Dynamics of Mitochondrial Metabolism. Cell 166, 1324–1337 e1311 (2016).

31. T. Kudo et al., Live-cell measurements of kinase activity in single cells using translocation reporters. Nat Protoc 13, 155–169 (2018).

32. J. R. Friedman et al., ER tubules mark sites of mitochondrial division. Science 334, 358–362 (2011).

33. J. Schindelin et al., Fiji: an open-source platform for biological-image analysis. Nat Methods 9, 676–682 (2012).

