## Supplemental Text and Figures for "Laterally transferred macrophage mitochondria act as a signaling source promoting cancer cell proliferation"

**This PDF file includes:**

Materials and Methods
Figs. S1 to S11
Caption for Movie S1

**Other Supplementary Materials for this manuscript include the following:**

Movie S1

### Materials and Methods

Cell culture of cell lines and peripheral blood mononuclear cells (PBMCs)

MDA-MB-231 and MCF10A cell lines were obtained through American Type Culture Collection and cultured according to their recommendations. MDA-MB-231 cells were cultured in 'DMEM complete media' containing: DMEM, high glucose (11965118, ThermoFisher) with 10% heat-inactivated fetal bovine serum (FBS; F4135, ThermoFisher). All cell lines were kept in culture for no more than 25 passages total.

*Genetic modification of PBMCs and differentiation into macrophages*

PBMCs were acquired from leukocyte filters obtained from de-identified blood donors (ARUP Blood Services). CD14+ monocytes were isolated from buffy coats and genetically modified with lentiviral vectors in the presence of virus-like particles packaging Vpx (to overcome restriction in myeloid cells) as previously described (*24*). Briefly, freshly harvested CD14+ monocytes were plated at a density of 4-5M cells per 10cm plate in 'macrophage culture media' containing: RPMI (11875119, ThermoFisher), 10% FBS (26140079, Thermo Fisher), 0.5% penicillin/streptomycin (P/S; P4333, Thermo Fisher), 10 mM HEPES (15630080, ThermoFisher), 0.1% 2-Mercaptoethanol (21985023, Thermo Fisher), recombinant human GM-CSF at 20 ng/ml (300-03, Peprotech) with the addition of polybrene (1 μg/ml), and supernatant containing Vpx particles (0.5 mL per 4M cells) to facilitate viral transduction. 30 mins after plating, 100 – 200 μL of concentrated lentiviral stock was added to the plated monocytes. 50% of the media was replaced on day 2 and a full media replacement occurred on day 4. Macrophages were used for experiments starting on day 6 or 7 after harvest of PBMCs unless otherwise noted.

*Generation of mito-FP and FP-TOMM20 stable cell lines*

We generated a modified pLKO.1 plasmid backbone with an accessible multiple cloning site (pLKO.1_MCS) for generation of mitochondrially targeted reporters. For mito-FP expression, we cloned the cytochrome oxidase subunit VIII mitochondrial targeting sequence and tagged it to *mEmerald* (referred to as mito-mEm) or *tagRFPt* (referred to as mito-RFP) and introduced these into the pLKO.1_MCS backbone in order to generate lentiviruses. pLKO.1 mito-mEmerald and pLKO.1 mito-TagRFP-T are available on Addgene (#174542 and 174543, respectively). For FP-TOMM20 expression, inserts containing the sequence of either *mEmerald* (mEmerald-TOMM20) or *mcherry* (mCherry-TOMM20) were fused to *TOMM20* and cloned into the pLKO.1 backbone and used to generate lentiviruses. For transduction, approximately 50,000 cells were plated into one well of a 6-well plate directly into the appropriate lentivirus supernant diluted 1:5 in DMEM complete media with a final concentration of 10 μg/mL polybrene (TR-1003-G, Sigma). After 48 - 72 hours, cells were expanded, and multiclonal populations were flow sorted for appropriate levels of fluorescent expression. All other transgenic cell lines were generated as outlined in subsequent sections.

mEmerald-TOMM20-N-10 (Addgene plasmid # 54282) and mCherry-TOMM20-N-10 (Addgene plasmid # 55146) were a gift from Michael Davidson.

Lentivirus production

pLKO.1_MCS plasmids containing the appropriate transgene were used to generate lentivirus as outlined in Johnson et al. 2020 (*24*). Briefly, 293FT cells in 15 cm plates were transfected with

PEI-max (24765, Polysciences) and plasmids for pCMV-VSV-G, psPax2, and transgene cassettes. The following day, cells were washed and cells were grown for an additional 36 hours in fresh media. Supernatants were harvested, passed through 0.45 μm syringe filters, and either used fresh or concentrated by ultracentrifugation as previously described (*24*). Lentiviral supernatants were used to transduce to cell lines as outlined in 'generation of mito-FP' section unless otherwise noted.

Flow cytometry

The following flow cytometry machines were used: a BD FACS Aria (equipped with 4 Lasers: 405, 488, 561, 640) referred to as Aria or a BD LSR Fortessa (5 Lasers: UV, 405, 488, 561, 640) referred to as the Fortessa. Technical details per experiment type are listed below.

*Stable line generation*

Cells were enzymatically dissociated using trypsin-EDTA (25200056, ThermoFisher) and resuspended in buffer consisting of 0.5% Bovine Serum Albumin (BSA; Sigma, A9418) in DPBS (14190250, ThermoFisher). Cells were sorted according to fluorescent intensity on the Aria and collected in the appropriate media containing 0.5% P/S.

*Mitochondrial transfer quantification*

For MDA-MB-231 and MCF10A cell lines: cells were enzymatically dissociated using trypsin-EDTA and stained as follows: cells were resuspended in 'staining buffer' (DPBS + 2% FBS) containing a human antibody against CD11b conjugated to the fluorophore Brilliant Violet 711 (BV711-CD11b; macrophage marker; Biolegend, 301344) was used at 1:20-40. After a 30 min incubation on ice, cells were washed and resuspended in cold DPBS for analysis on the Fortessa. The background level of mEmerald fluorescence was set at 0.2% based on a fully stained monoculture control. This gate was defined by FACS-isolating co-cultures of mito-RFP MDA-MB-231/mito-mEm macrophages and determining a gate that accurately isolated MDA-MB-231 cells containing macrophage mitochondria.

*Mitochondrial transfer quantification of PDxO containing co-cultures*

Hanging drop co-cultures suspended in Growth Factor Reduced Matrigel (354230, Corning) were pooled and dissociated using a solution of Dispase II (50U/mL; 17105041, Fisher Scientific) followed by TrypLE Express (12605010, Thermo Fisher). Cells were then incubated in TrueStain FcX (422301, ThermoFisher) at 1:33 diluted into staining buffer for 10 minutes at room temperature. Primary human antibodies against CD326 conjugated to PE (PE-EpCam; PDxO marker; 369806, Biolegend) and BV711-CD11b were added at 1:20 and 1:40, respectively. After 30 mins on ice, cells were washed and resuspended in cold DPBS for analysis on the Fortessa. The background level of mEmerald fluorescence in the 'transfer gate' was set at 0.2% based on a fully stained monoculture control.

*Quantification of Ki67 and DNA content*

Co-cultures were enzymatically dissociated with trypsin-EDTA and incubated in staining buffer containing anti-human BV711-CD11b at 1:40 for 30 minutes on ice. Cells were then fixed and stained using the eBioscience Foxp3/Transcription Factor Staining Buffer Set (00-5523-00, ThermoFisher) and according to manufacturer's instructions. Cell were stained with an APC conjugated Ki67 antibody (17-5699-42, ThermoFisher) at 1:20 for 30 minutes followed by a

3μM DAPI solution for 10 minutes. Cells were resuspended in cold DPBS for analysis on the Fortessa. The background level of mEmerald fluorescence in the 'transfer gate' was set at 0-0.2% based on a fully stained monoculture control.

Single-Cell RNA-seq

From FACS-isolated populations, a cDNA library was generated using the 10X genomics Single Cell 3' Gene Expression Library V3 and amplified according to the manufacturer's protocol. The resulting libraries were sequenced on a NovaSeq 6000 resulting in approximately 100K mean reads per cell. The raw sequencing data were processed using CellRanger 3.02 (*https://support.10xgenomics.com/*) to generate FASTQ files, aligned to GRCh38 (Ensemble 93), and a gene expression matrix for individual cells based on the unique molecular indices was generated. The resultant filtered gene-cell barcode matrix was imported into SEURAT version 4 (*25*) with R studio version 1.3.1093 and R version 4.03. We first performed quality control by determining the mean and standard deviation of genes per cell and filtered out all cells that were more than 1.5 standard deviations away from the mean. The reads were then scaled and normalized using SEURAT 'sctransform' function (*26*). Using the normalized data, we determined differential gene expression in the MDA-MB-231 population that received macrophage mitochondria compared to those that did not, using a non-parametric Wilcoxon rank sum test with the SEURAT 'FindMarkers' function. Lastly, the differential expression data were exported from R and pathway enrichment analysis was performed using Qiagen's Ingenuity Pathway Analysis software (*27*). Single-cell RNA-sequencing data are available with GEO accession number GSE181410. The analysis code for single-cell RNA-sequencing analysis is available on GitHub (https://github.com/rohjohnson-lab/kidwell_casalini_2021).

Trans-well experiments

Approximately 40,000 mito-RFP MDA-MB-231 and 80,000 mito-mEm macrophages were plated in trans-wells (3401, Corning) in the conditions listed in Fig. S1E. Cells were analyzed after 24 hours with flow cytometry as indicated in 'mitochondrial transfer quantification' section.

Live cell imaging of co-cultures with cell-permeable dyes

Imaging was performed using either a Zeiss LSM 880 with AiryScan technology (Carl Zeiss, Germany) and a 63x/1.4 NA oil objective or a Leica Yokogawa CSU-W1 spinning disc confocal microscope with a Leica Plan-Apochromat 63x/1.4 NA oil objective and iXon Life 888 EMCCD camera. Images taken on the LSM 880 were acquired using the AiryScan Fast mode. For all live imaging, cells were maintained at 37ºC, 5% $CO_2$ with an on-stage incubator.
MDA-MB-231 cells and primary macrophages stably expressing the appropriate transgenes were mixed in a 1:2 ratio and plated at an approximate density of 300,000 cells directly onto 35mm glass bottom dishes (FD35-100, World Precision Instruments) for all live imaging experiments unless otherwise noted. Duration of co-culture is indicated in main text or figure legend.

For detection of mitochondrial DNA and the nucleus, Hoechst 33342 (B2261, Sigma) was diluted into the culture media to a final concentration of 5 μg/mL. After 10 mins, cells were washed, and complete media was replaced before imaging.

For detection of mitochondrial membrane potential with Mitotracker Deep Red (MTDR; M22426, ThermoFisher) was diluted into serum-free DMEM media (11965118, ThermoFisher) at a final concentration of 25 nM and incubated at 37°C for 30 mins. Following incubation, cells were washed, and complete media was replaced before imaging.

For detection of mitochondrial membrane potential with Tetramethylrhodamine, Methyl Ester, Perchlorate (TMRM; T668, ThermoFisher) was diluted into serum-free DMEM media at a final concentration of 100 nM and incubated at 37°C for 30 mins. Following incubation, cells were washed, and complete media was replaced before imaging.

For detection of acidic vesicles, Lysotracker Blue (L7525, ThermoFisher) was diluted to a final concentration of 75 nM in serum-free DMEM media and incubated at 37°C for 30 mins. Following incubation, cells were washed and complete media was replaced before imaging.

For detection of plasma/vesicular membranes, MemBrite 640/660 (Biotium, 30097) was used final concentration of 1:1000 and stained according to manufacturer's instructions.

For detection of ROS, Carboxy-$H_2$DCFDA (C400, ThermoFisher) was diluted to 5 μM into warmed HBSS (14025092, ThermoFisher) and incubated at 37ºC for 15-30mins. After incubation, cells were washed with HBSS and warmed complete media was replaced before imaging.

Live imaging of sorted recipient cells

MDA-MB-231 cells were harvested and stained as indicated in 'mitochondrial transfer' section of flow cytometry methods. Cells were sorted on the Aria directly into media containing 0.5% P/S. Sorted cells were plated directly onto imaging dishes coated with CellTak (354240, Corning) and allowed to attach at 37ºC for up to 4 hours before staining and live imaging.

Quantitative phase imaging (QPI)

Mito-RFP MDA-MB-231 cells and mito-mEm macrophages were seeded in a 1:2 ratio at a density between 90,000 - 120,000. They were plated directly onto imaging dishes 24 hours prior to the start of imaging. QPI images were acquired on Olympus IX83 inverted microscope (Olympus Corporation, Japan) with Phasics SID4 camera (Phasics, France) and Thorlabs 623 nm wavelength DC2200 LED (Thorlabs, USA). The microscope was operated in brightfield with Olympus UPLFLN 40X objective and a 1.2X magnifier in front of camera, giving 48X magnification. Fluorescence images were acquired using X-Cite 120LED illumination (Excelitas technologies, USA) and an R1 Retiga camera (Cairn research Ltd, UK) with GFP (Olympus Corporation U-FBNA) and RFP (IDEX health & science, USA mCherry-B-000) filter cubes. Cells were maintained at 37ºC temperature, 5% $CO_2$ and 90% humidity with an Okolab (Okolab, Italy) on-stage incubator on a Prior III Proscan microscope stage (Prior Scientific Instruments Ltd., UK). Automation was performed with MicroManager open-source microscopy software via MATLAB 2012b. QPI images of 40 positions per imaging set, four replicate (biological replicate) imaging sets total, were acquired every 15 minutes with fluorescence images acquired in an alternate subset of locations every 15 minutes for 48 hours to reduce phototoxicity.

*QPI data analysis*

QPI and fluorescent images were analyzed with MATLAB 2019a. Cell phase shift images were background corrected using sixth order polynomial surface fitting, and converted to dry mass ($m$)

map, using, $m = \int \frac{1}{\alpha} \phi \lambda dA$, where $\lambda$, is the wavelength of source light = 0.623 μm, $\alpha$, specific refractive increment = 0.185 $\mu m^3/pg$, $A$, image pixel area = 0.36 $\mu m^2$/pixel, and $\phi$ is the phase shift in fraction of a wavelength at each pixel. Cell dry mass maps were then segmented using a Sobel filter for edge detection and tracked over time (*28*). Specific growth rate of each tracked cell was computed as the slope of a linear, least-squares best fit line to mass over time data normalized by cell average mass. Fluorescent mitochondria images were resized to match QPI images, and overlayed with corresponding QPI image segmentation mask to measure the integrated fluorescence intensity of every cell, normalized by cell area. Macrophage mitochondria high frequency punctae signal in MDA-MB-2321 cells were separated from the high intensity, low spatial frequency of the macrophage mitochondria network fluorescence signal using the rolling ball filter in MATLAB. The size of the rolling ball was 4.8 to 9 μm, chosen to be just above the average size of mitochondrial punctae based on the quantity of mitochondria transferred and retained in the MDA-MB-231 cells. Cells with RFP signal 1.5 times more than the background were identified as MDA-MB-231 cells, and with mEmerald fluorescent signal double that of background as macrophages. MDA-MB-231 cell tracks were then binned based on the presence or absence of mEmerald+ mitochondrial punctae, indicating transfer from macrophages. The specific growth rate of each cell was calculated as the slope of a least-squares linear fit to QPI mass vs time data divided by the average mass of the cell. The code for automated tracking of cell mass from QPI and fluorescence data and computing growth rates of the different groups of cells is available on GitHub (https://github.com/Zangle-Lab/Macrophage_tumor_mito_transfer).

*Cytokinesis analysis*
Cytokinesis rate was calculated by tracking cells manually to confirm division of cells in less than the maximum doubling time expected (40 hours). Cells leaving the imaging frame in less than 30 hours were omitted from the cytokinesis calculation.

*Lineage analysis*
The average specific growth rate of MDA-MB-231 parent and daughter cell was calculated by manually annotating mass versus time tracks from mass tracking based on the presence of mEmerald+ punctae. The difference in growth of daughter cells that did or did not inherit mitochondria from mitochondria containing parents was observed by normalizing the mass of each daughter by its initial mass at birth.

ROS biosensor line generation, imaging, and quantification
MDA-MB-231 cells were transfected with the following plasmids: pLPCX mito-Grx1-roGFP2 (*10*) and pLPCX mito-roGFP2-Orp1 (*11*) using the Polyplus-transfection jetPRIME DNA/siRNA transfection kit (55-131, Genesee Scientific) according to the manufacturer's instructions. Cells were allowed to recover for 3-7 days and then sorted for expression. Cells were passaged every 3-5 days and sorted as needed to maintain a high percentage of expressing cells. Biosensor-expressing MDA-MB-231 lines were co-cultured with mito-RFP expressing macrophages for 24 or 48 hours and imaged on the Zeiss LSM 880. Cells were sequentially imaged (per z-plane) for the presence of transferred macrophage mitochondrial (Ex. 561nm, Em. BP 570-620nm + LP 645nm) and the biosensor in its a reduced (Ex. 488nm, Em. BP 420-480nm + BP 495-550nm) and oxidized (Ex. 405nm, Em. BP 420-480nm + BP 495-550nm) form.

Images were initially processed using Zen software (see image analysis section) and further analysis was performed using FIJI as indicated in Morgan et al. 2011 (*29*).

pLPCX mito Grx1-roGFP2 (Addgene plasmid # 64977) and pLPCX mito roGFP2-Orp1 (Addgene plasmid #64992) were a gift from Tobias Dick.

Mito-KillerRed line generation and imaging

To generate *3xHA-killerred-OMP25,* a plasmid containing *3xHA-EGFP-OMP25 (30)* was used as a template and the sequence of KillerRed replaced EGFP. The entire transgene was then cloned into the pLKO.1 backbone. pLKO.1 3xHA-KillerRed-OMP25 is available on Addgene (#174544). MDA-MB-231 cells were transduced with lentiviral supernant that packaged the *3xHA-killerred-OMP25* transgene, allowed to recover and were cell sorted to select for the appropriate level of fluorescent expression. Cells expressing both mito-mEm and mito-KillerRed were generated in parallel to confirm the correct localization of the mito-KillerRed (data not shown).

pMXs-3XHA-EGFP-OMP25 was a gift from David Sabatini (Addgene plasmid #83356).

*For generation of mt-ROS with the mito-KillerRed cell line*

MDA-MB-231 mito-KillerRed-expressing cells were labelled with Carboxy-$H_2$DCFDA as described above. Using a Leica Yokogawa CSU-W1 spinning disc confocal microscope equipped with a 2D-VisiFRAP Galvo System Multi-Point FRAP/Photoactivation module, MDA-MB-231 mito-KillerRed-expressing cells were imaged at 488nm (for DCFDA detection) and 561nm (for mito-KillerRed detection) at a time interval of 2 seconds. After 2 frames, a ~2um x 2um region of interest (ROI) of mito-KillerRed was photobleached using a 561 laser (100% laser power, 5ms, 1 cycle), and continuous imaging at 488nm and 561nm allowed for DCFDA quantification and mito-KillerRed photobleaching, respectively.

*To quantify cell division upon ROS production*

Cells were stained with 5 μg/mL Hoescht 33342 as described above to visualize nuclei. Multiple stage positions were established such that control experiments, in which a cytoplasmic ROI without mito-KillerRed expression that was photobleached using identical parameters, as well as a no-photobleaching control, could be imaged simultaneously with experimental photobleached cells. Approximately 8-10 cells of each category – photobleached in mito-KillerRed-expressing regions, photobleached in control cytoplasmic non-expressing regions, or not photobleached – were imaged by acquiring Z-stacks (1 μm step size) every 15 minutes for 18 hours. Cell division was quantified by visualizing nuclear division with FIJI software.

ERK-KTR generation

MDA-MB-231 cells were transduced with lentiviral supernant that was packaged using either pLentiPGK Blast DEST ERKKTRmRuby2 or pLentiPGK Puro DEST ERKKTRClover plasmids (*31*) as outlined in 'generation of mito-FP' section. Cells were allowed to recover post-infection, sorted for fluorescent expression, and maintained as stable cell lines.

pLentiPGK Blast DEST ERKKTRmRuby2 (Addgene plasmid # 90231) and pLentiPGK Puro DEST ERKKTRClover (Addgene plasmid # 90227) were a gift from Markus Covert.

ERK-KTR-mClover and mito-KillerRed generation and imaging:

A stable MDA-MB-231 line expressing mito-KillerRed was transduced with lentivirus that was packaged using a pLentiPGK Puro DEST ERKKTRClover plasmid. Cells were sorted for expression of both mito-KillerRed and ERK-KTR-mClover and maintained as a stable line.

*For mt-ROS generation and ERK-KTR imaging*

MDA-MB-231 cells expressing mito-KillerRed and ERK-KTR-mClover were stained with 5μg/mL Hoescht 33342 as described above to visualize nuclei. Using a Leica Yokogawa CSU-W1 spinning disc confocal microscope equipped with a 2D-VisiFRAP Galvo System Multi-Point FRAP/Photoactivation module, MDA-MB-231 cells expressing mito-KillerRed and ERK-KTR-mClover were imaged every 1 minute with 561nm (for mito-KillerRed) and 488nm (for ERK-KTR-mClover) and 405nm (for nuclei) lasers. A ~2 um x 2 um ROI of KillerRed+ mitochondria was photobleached using a 561 laser (100% laser power, 5ms, 1 cycle), and continuous imaging at 488nm and 561nm allowed for visualization of ERK-KTR-mClover translocation and mito-KillerRed photobleaching, respectively. Multiple stage positions were set such that control experiments, in which a cytoplasmic region without mito-KillerRed expression that was photobleached using identical parameters, could be imaged simultaneously with experimental photobleached cells.

*ERK-KTR quantification with FIJI*

ERK-KTR-mClover translocation was quantified every 10 minutes by taking maximum projections of Z-planes only encompassing the cell nucleus. Using FIJI software, a ROI was drawn in the nucleus guided by the Hoescht staining, and the MFI of ERK-KTR-mClover was quantified in this region. The same ROI was moved outside of the nucleus to a cytoplasmic region devoid of mitochondria, and the MFI of ERK-KTR-mClover was quantified. This analysis was performed for each timepoint after photobleaching. The values are then used to calculate a cytoplasmic:nuclear ratio at each time point, and normalized to 1 at time point zero.

Quantification of ERK-KTR using the Amnis ImageStream

To quantify translocation of the ERK-KTR-mRuby we used the Amnis Imagestream mk II with ISX software (version 201.1.0.725). Mito-mEm macrophages were co-cultured with ERK-KTR-mRuby+ MDA-MB-231 cells for 24 hours. Samples were prepared as indicated in ‘quantification of Ki67 and DNA content’ section with the exception that we did not stain for intracellular markers. Images were captured with the 40x objective and sample collect flow was set to low, as this allows for higher image resolution. Using Image Data Exploration and Analyses Software (IDEAS; version 6), we quantified translocation using two metrics: 1) the IDEAS translocation Wizard and 2) custom-generated program to detect cytoplasmic (cyto) and nuclear (nuc) ERK-KTR intensities to calculate a cyto:nuc ratio (as indicated in Fig. S6B-S7B). The translocation wizard is a pre-built program made to detect the nuclear translocation of a probe. It does this by making a pixel-by-pixel correlation between the probe of interest (ERK-KTR) and the nuclear image (DAPI). The program gives each cell a score indicating how similar the two fluorescent images are. A high score suggests the images are similar (more nuclear translocation) and a low score suggests that the images are less similar (less nuclear translocation). We also quantified ERK-KTR translocation by generating a custom masking strategy to quantify the mRuby median fluorescent intensity (MFI) in the cytoplasm and nucleus using IDEAS software. To identify nuclear mRuby, we manually set a threshold of DAPI signal and reported the mRuby MFI of pixels within that threshold range. To quantify the cytoplasmic mRuby fraction we reported the mRuby MFI from outside the threshold. These values are then used to calculate a cyto:nuc ratio.

<u>ERKi and PMA Drug treatments</u>
SCH772984 (ERKi; 7101, SelleckChem) and Phorbol 12-myristate 13-acetate (PMA; S7791, Selleckchem) was dissolved in 100% DMSO to make 10 mM stock solutions and stored at -80°C. No individual aliquot went through more than 2 freeze-thaw cycles. The stock solution was thawed and then diluted directly into complete media for a final concentration of 1 μM for ERKi and 100 nM for PMA. For all ERKi experiments, co-cultures were treated at the time of plating and for a duration of 24 hours. All PMA treatments, plated cells were treated for 1 hour prior to harvest and analysis.

<u>Macrophage activation and verification</u>
For macrophage activation, macrophages were harvested and differentiated as indicated in 'cell culture of PBMCs' section. Between days 6-7 of differentiation, IFN-γ (3000-02, Peprotech, 20 ng/mL) for M1 activation or IL-4 + IL-13 (200-04, 200-13, Peprotech, 20 ng/mL) for M2 activation were added to culture media for 48 hours before experiments were conducted. To confirm M1 and M2 activation, macrophages were collected and stained for known surface markers for M1 (CD86; 62-0869-42, Thermofisher) and M2 (CD206; 321110, Biolegend) activation. Flow cytometry was performed on the Fortessa to observe changes in fluorescent intensities across M0, M1, and M2 macrophages populations (Fig. S10A).

<u>Immunofluorescence and analysis of mitochondrial morphology</u>
Mito-mEm expressing macrophages were co-cultured with mito-RFP+ MDA-MB-231 cells for 24 hours and fixed with warm 4% PFA with 5% sucrose in 1x DPBS for 20 minutes and permeabilized with 0.2% Triton X-100 in 1x DPBS (9002-93-1, Sigma). Cells were stained with chicken α-GFP (AB13970, Abcam) and Rabbit α-RFP (AB62341, Abcam) antibodies at 1:500 and 1:1000, respectively. The following secondary antibodies were used: Alexa Fluor 488 AffiniPure Goat anti-Chicken (103-545-155, Jackson ImmunoResearch) and IgG (H+L) Cross-Adsorbed Goat anti-Rabbit Alexa Fluor 555 (A21428, Invitrogen) both at 1:500. Cells were subsequently stained with 1 μg/mL DAPI (D9542, Sigma) in DPBS for 10 minutes. Cells were then mounted with ProLong Diamond Antifade Mountant (P36965, ThermoFisher) and stored at 4°C before imaging. Imaging was performed using the Zeiss LSM 880 using the AiryScan fast mode. AiryScan processed images (see image analysis section) were used to quantify mitochondrial morphologies with the FIJI plug-in, Mitochondrial Network Analyses (MiNA; Fig. S10B-C). Pre-processing parameters: Manually select top and bottom of the cell of interest, exclude any space above and below the cell as this can introduce background noise. 3D project cell. Unmask sharp Radius (5), Mask Weight (0.6), Median 3D (0.5, 0.5, 0.5), Make binary (Otsu), Skeletonize, Analyze skeleton 2D/3D. A 'mitochondrial fragment' was defined as a mitochondrion with 0-1 branches, 0 junctions, and a length between 0-2 μm.

<u>DRP1 knockdown and overexpression</u>
Monocytes were isolated as indicated in 'cell culture of PBMCs' section and transduced with lentiviruses to express mito-mEm and either non-target (nt) short hairpin (sh) RNA (SHC002,

Sigma), *DRP1*-shRNA (TRCN0000001097, Sigma; gene target HGNC ID 2973) or mCherry-DRP1 (*32*). All constructs were either produced or cloned into the pLKO.1 backbone.

mCh-Drp1 (Addgene plasmid # 49152) was a gift from Gia Voeltz.

rt-qPCR verification of genetic knockdown

RNA from nt-shRNA and *DRP1*-shRNA expressing macrophages were isolated from 3 independent macrophage donors. To isolate RNA we used standard TRIzol/chloroform RNA isolation techniques. cDNA libraries were made using SuperScript III Reverse Transcriptase (18080093, ThermoFisher), according to manufacturer's instructions. DRP1-knockdown was verified via qRT-PCR with Power SYBR Green Mast Mix (4368511, ThermoFisher). Primers were designed with NCBI primer design, commercially produced by Integrated DNA Technologies and tested for specificity with standard PCR. Primers were as follows; DRP1-F: AGAAAATGGGGTGGAAGCAGA, DRP1-R: AAGTGCCTCTGATGTTGCCA, GAPDH-F: AGCCACATCGCTCAGACA, GAPDH-R: ACATGTAAACCATGTAGTTGAGGT. Cycle Thresholds (CT) values were determined by averaging 3 technical replicates from 3 biological samples. Control ΔCT: expression was normalized to *GAPDH* by subtracting the *DRP1* CT value of the nt-shRNA expressing macrophages from the *GAPDH* CT value of the same sample. Target gene, *DRP1* ΔCT: *DRP1* CT values of the *DRP1*-shRNA expressing macrophages were subtracted from the *GAPDH* CT values of the same sample. The ΔΔCT values was calculated by subtracting *DRP1* ΔCT – control ΔCT. Normalized target gene expression was calculated ($2^{-\Delta\Delta CT}$) and used to determine % knockdown (($1-2^{-\Delta\Delta CT}$)*100).

PDxO culture and co-culture with macrophages

PDxO cell lines HCI-037 and HCI-038 were generated and maintained as described in Gullien et al. 2021 (*19*). Like MDA-MB-231 cells, these models are estrogen and progesterone receptor negative and HER2 negative (triple negative breast cancer). For co-culture with macrophages, mature PDxOs were dissociated from Matrigel with a Dispase II solution followed by treatment with TrypLE Express to generate a suspension of single cells. PDxO cells were then mixed with mito-mEm macrophages (differentiated for 6-8 days) in a 1:2 ratio at a density of 90,000 cells total per hanging drop culture. Macrophage media was used for hanging drops (for media components, see isolation of PBMCs section) and they were suspended from the lid of a tissue culture plate to allow for cell aggregation for 24 hours and then pooled and embedded into Growth Factor Reduced Matrigel. Embedded hanging drop cultures were then allowed to incubate for 72 hours and were then analyzed for mitochondrial transfer with flow cytometry (see flow cytometry section).

Image analysis

All images taken with the Airyscan detector on the Zeiss LSM 880 were subjected to deconvolution using the Zen software (Carl Zeiss) with 'auto' settings (referred to as AiryScan processed). Maximum intensity projections of selected z-planes were generated using Zen or FIJI software (*33*). Linear adjustments to the brightness and contrast were made using FIJI. Images were cropped and panels were assembled using Adobe Photoshop and Illustrator, respectively (Adobe, Inc.).

Graphical representations and statistical analysis

All graphs were generated using Prism software (v9, GraphPad). Statistical analyses were performed using both Excel (v16.51, Microsoft) and Prism. Statistical tests used and p-value ranges are indicated in each figure legend. Flow cytometry data and representations were analyzed and generated using FlowJo software (v10.7, BD).

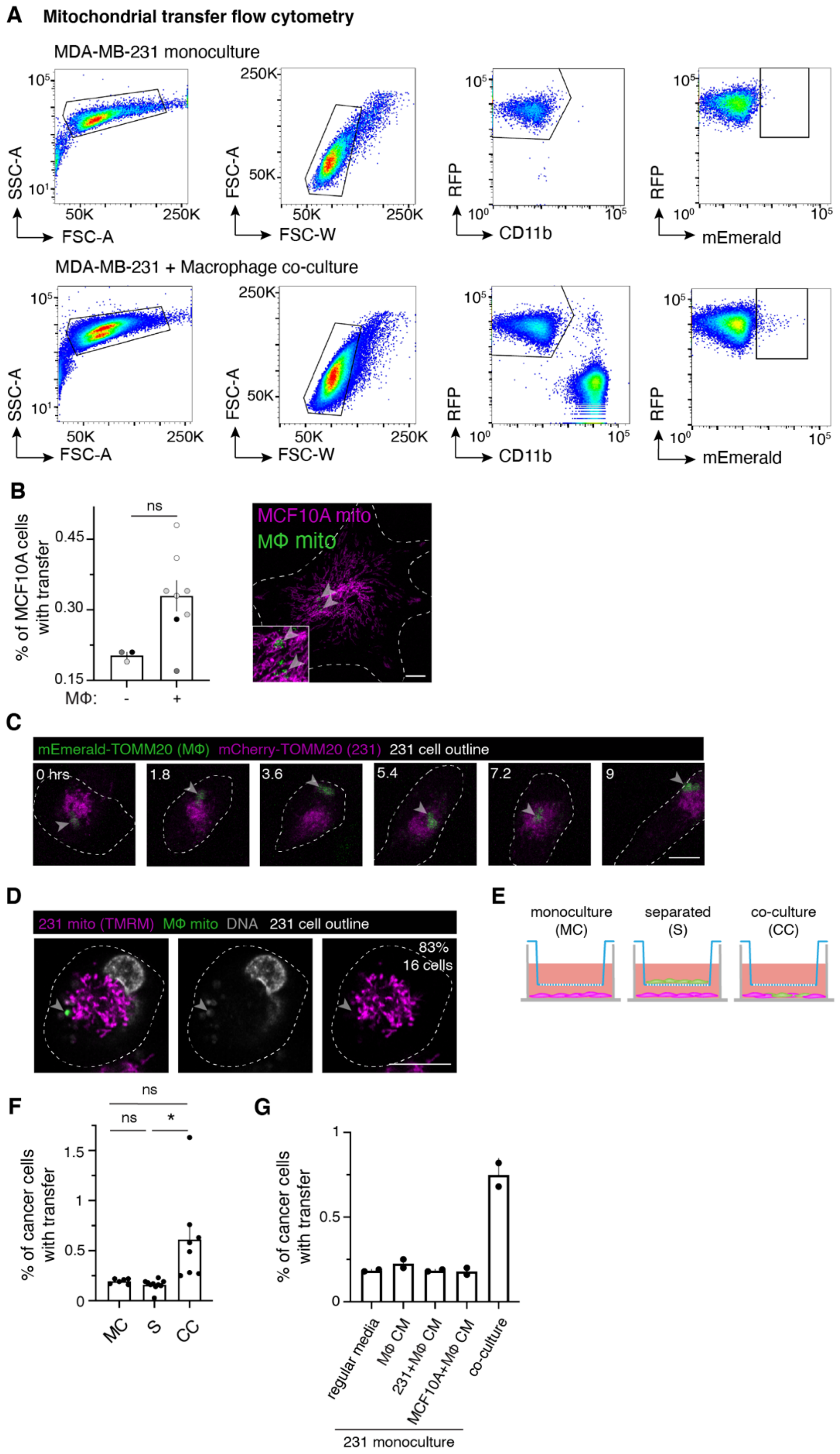
A Mitochondrial transfer flow cytometry
MDA-MB-231 monoculture
SSC-A
FSC-A
FSC-W
RFP
CD11b
mEmerald
MDA-MB-231 + Macrophage co-culture
B
% of MCF10A cells with transfer
ns
MΦ: - +
MCF10A mito
MΦ mito
C
mEmerald-TOMM20 (MΦ) mCherry-TOMM20 (231) 231 cell outline
0 hrs
1.8
3.6
5.4
7.2
9
D
231 mito (TMRM) MΦ mito DNA 231 cell outline
83%
16 cells
E
monoculture (MC)
separated (S)
co-culture (CC)
F
% of cancer cells with transfer
ns
*
MC
S
CC
G
regular media
MΦ CM
231+MΦ CM
MCF10A+MΦ CM
co-culture
231 monoculture

**Fig. S1. Macrophages transfer mitochondria to cancer cells.**

(A) Representative flow cytometry plots of mito-RFP MDA-MB-231 monocultured cells used as a control (top) and mito-RFP 231/mito-mEm macrophage co-cultures after 24 hours (bottom). (B) Left: Bar graph quantifying mitochondrial transfer to MCF10A cells after 24 hours. (N= 3 donors indicated by color). Right: Confocal image showing macrophage mitochondria (green, arrowhead) in a MCF10A cell (magenta). (C) Stills from a time-lapse showing a 231 cell (mCherry-TOMM20, magenta) containing transferred macrophage mitochondria (mEmerald-TOMM20, green, arrowhead). Timepoints indicated in upper left. (D) Single Z-plane of a 231 cell labeled with mitochondrial dye TMRM (magenta) with macrophage mitochondria (green, arrowhead) containing DNA (gray). 83% of transferred mitochondria contain DNA (N= 16 cells). (E) Schematic depicting trans-well experiments. 231 cells (magenta) were plated alone (left), separated from macrophages (green; middle), or plated together (right). (F) Percent of cancer cells with transfer in conditions depicted in (E) (N= 3 donors). (G) Conditioned media (CM) experiments showing percent of cancer cells with transfer when co-cultured in media type listed on the x-axis. Each dot represents one replicate for all panels. Error bars represent standard error of the mean (SEM) and scale bars are 10 µm. Mann-Whitney test (B), ANOVA 2-way (F), $^{*}p<0.05$; $^{**}p<0.01$.

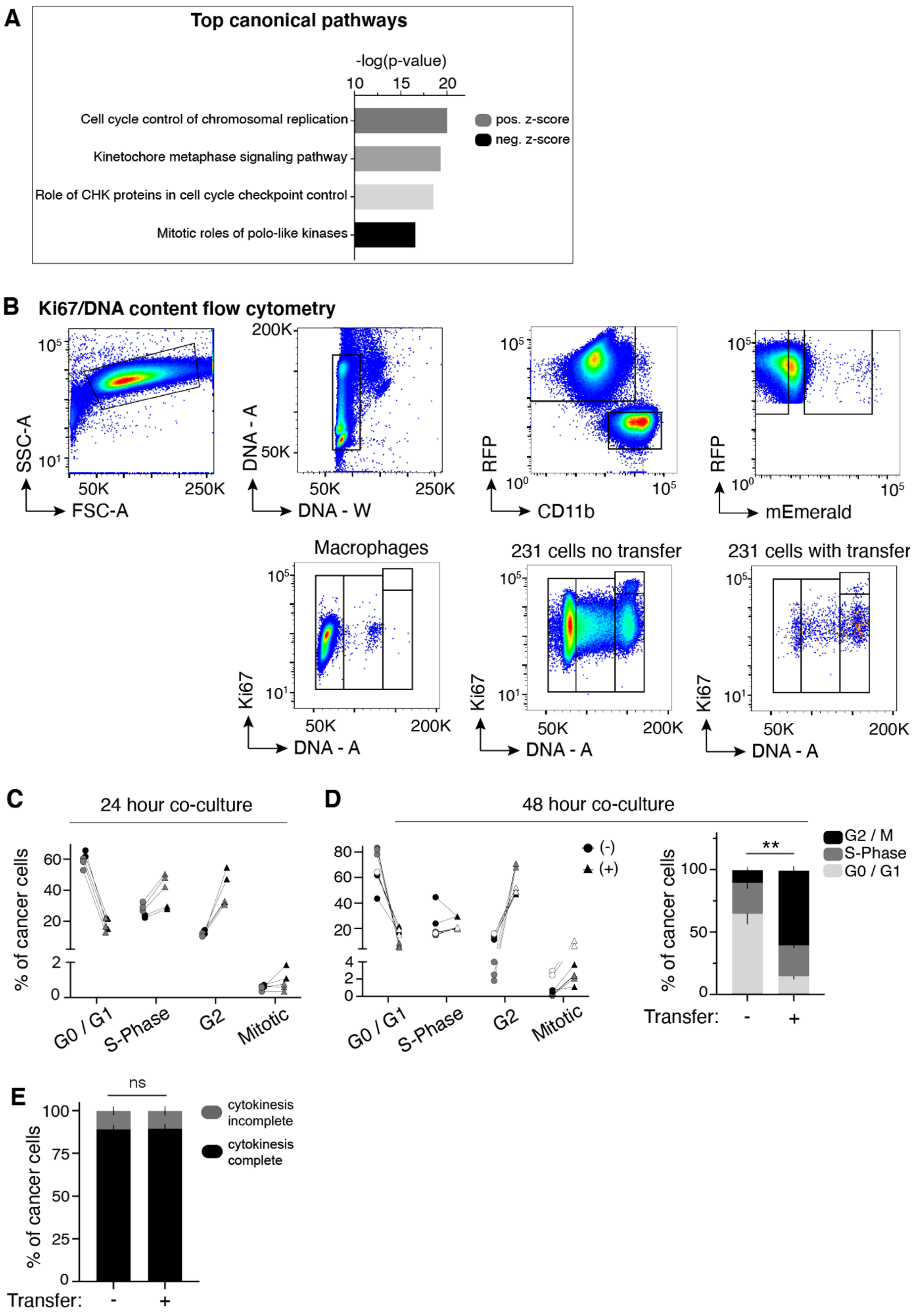
A
Top canonical pathways
-log(p-value)
10
15
20
Cell cycle control of chromosomal replication
Kinetochore metaphase signaling pathway
Role of CHK proteins in cell cycle checkpoint control
Mitotic roles of polo-like kinases
pos. z-score
neg. z-score
B
Ki67/DNA content flow cytometry
SSC-A
FSC-A
DNA - A
DNA - W
RFP
CD11b
RFP
mEmerald
Macrophages
231 cells no transfer
231 cells with transfer
Ki67
DNA - A
C
24 hour co-culture
% of cancer cells
G0 / G1
S-Phase
G2
Mitotic
D
48 hour co-culture
(-)
(+)
G2 / M
S-Phase
G0 / G1
Transfer:
E
ns
% of cancer cells
cytokinesis incomplete
cytokinesis complete
Transfer:

**Fig. S2: Cancer cells will macrophage mitochondria exhibit increased proliferation.**

(A) Ingenuity Pathway Analysis of RNA-seq data reveals significant changes in canonical cell proliferation pathways in co-cultured cancer cells that received macrophage mitochondria compared to co-cultured cancer cells that did not. (B) Representative flow cytometry plots of mito-RFP 231/mito-mEm macrophage co-cultures stained for Ki67 and DNA content. (C) 24 hour data from Fig 1E but separated as percent of cancer cells in each phase of the cell cycle with (triangle) or without transfer (circle). (D) 48 hours of co-culture as described in (C) with aggregate data in stacked bar graphs to the right. Each pair in C and D represents cells within one technical replicate and color indicates matched replicates from one experiment (N= 2 donors for 24 hour data and 3 donors for 48 hour data; statistics displayed for G2/M only). (E) Time-lapse imaging was used to quantify the amount of co-cultured 231 cells that did (black) or did not complete (gray) cytokinesis over a 48 hour period (N= 416 cells, 4 donors). Error bars represent SEM. Unpaired t-test with Welch's correction (D, G), Mann-Whitney test (E).

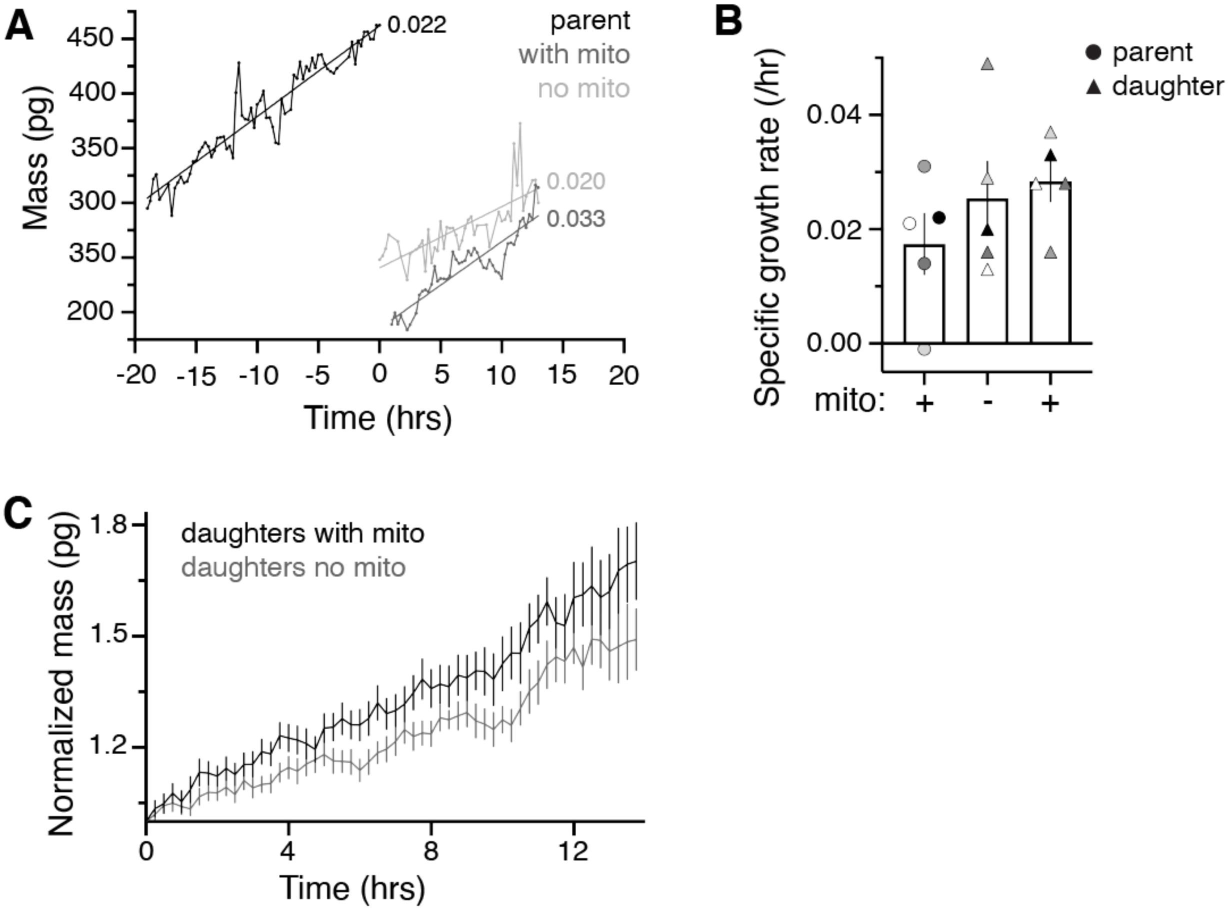


**Fig. S3. Mitochondrial transfer leads to sustained increased growth rate in daughter cancer cells.**
(A) QPI data of mass over time measurements for a single triad consisting of one parent cell (black) and two daughters that inherited (gray) or did not inherit (light gray) the parent's macrophage mitochondria. Specific growth rate (slope of best fit line normalized by average mass) is listed next to line trace of corresponding cell. (B) QPI data of specific growth rate of 5 individual triads (indicated by color). Parent and daughter cells are indicated with shape (legend on graph). (C) QPI data of normalized mass in picograms (pg) over time in hours (hrs) for daughter cells that did (black) or did not (gray) inherit the parent's macrophage mitochondria normalized to daughter cell initial mass. Error bars represent SEM.

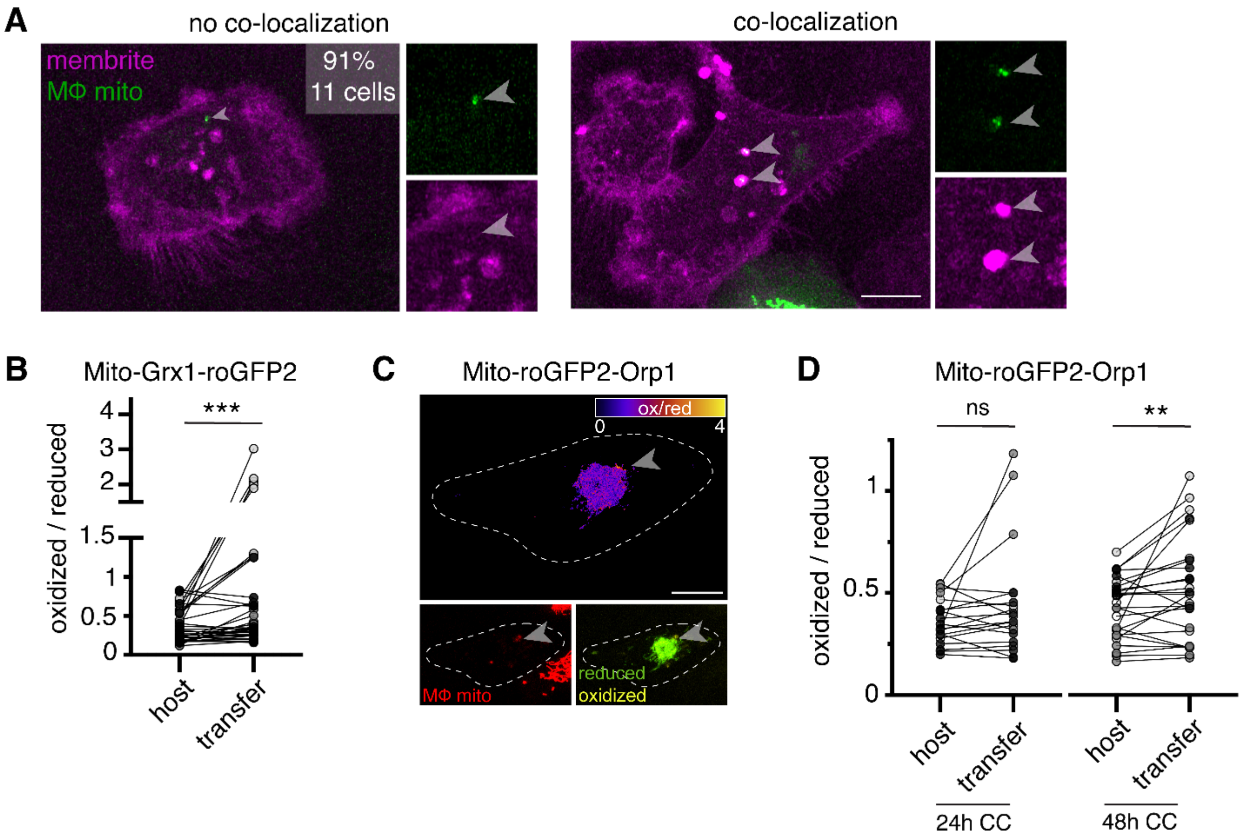


**Fig. S4. Transferred mitochondria accumulate reactive oxygen species.**

(A) Recipient 231 cells were stained with a dye, MemBrite, that marks both the plasma and vesicular membranes. 91% of transferred mitochondria (green, arrowheads) did not co-localize with MemBrite signal (magenta, left panel) whereas 9% did (right panel) (N= 11 cells, 1 donor). (B) Ratiometric measurements of the mito-Grx1-roGFP2 sensor per 231 cell (paired dots) at a region of interest (ROI) containing the recipient host mitochondrial network (host) or a transferred mitochondria (transfer). Cells were co-cultured with macrophages for 48 hours (N=37 cells, 3 donors). (C) Ratiometric quantification of mito-roGFP2-Orp1 biosensor mapped onto recipient 231 cell in the fire LUT (top). Bottom left panel shows macrophage mitochondria (bottom left, red, arrowhead) and bottom right shows mito-roGFP2-Orp1 (green and yellow). (D) Ratiometric measurements of the mito-roGFP2-Orp1 sensor per 231 cell (paired dots) at a ROI containing the host mitochondrial network (host) or a transferred mitochondria (transfer). Cells were co-cultured with macrophages for 24 (N=21 cells, 3 donors) or 48 hours (N=26 cells, 3 donors). For all panels, individual donors are indicated as shades of gray and scale bars are 10 µm. Wilcoxon matched-pairs signed rank test, $**p<0.01$; $***p<0.0001$.

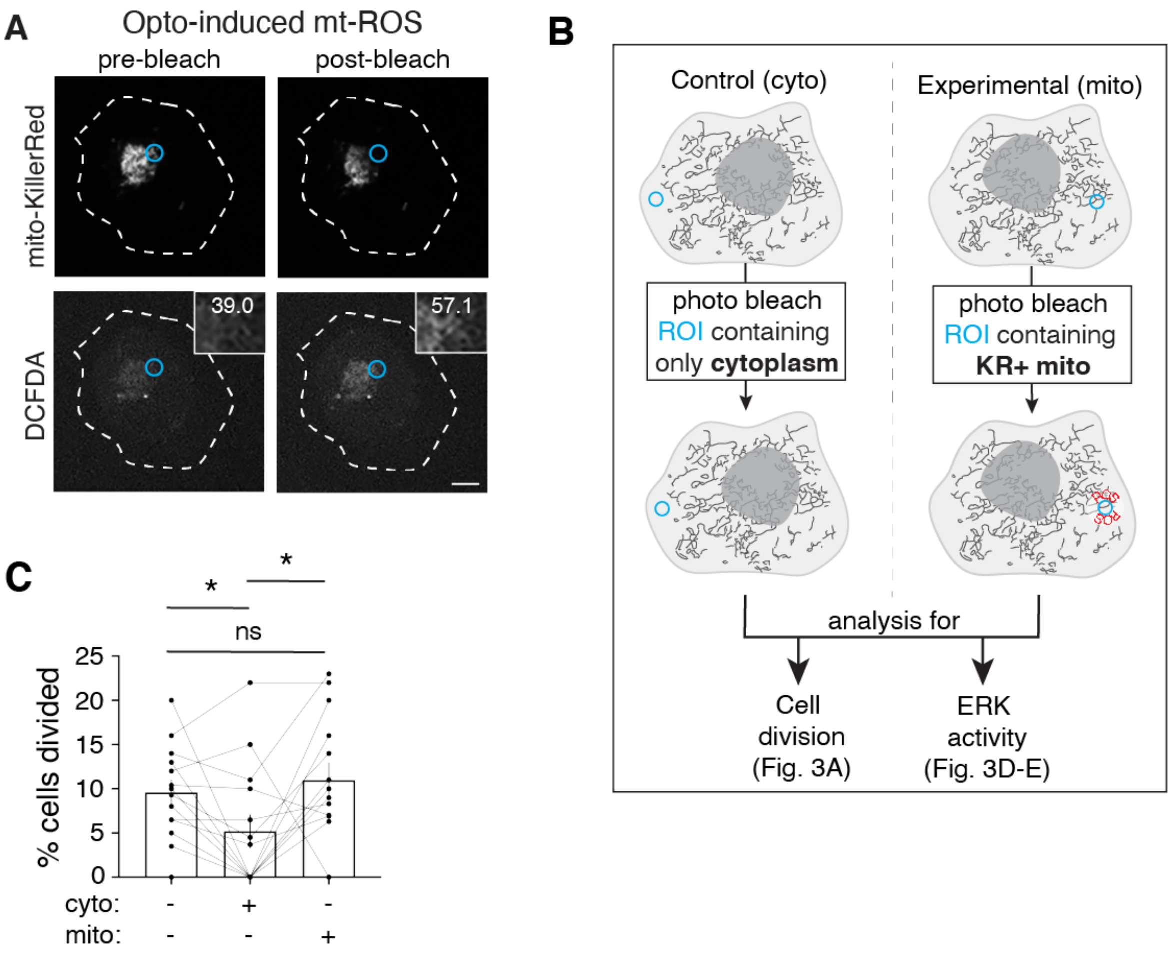


**Fig. S5. Inducing reactive oxygen species results in cancer cell proliferation.**

(A) Photobleaching a region of interest (ROI; blue) containing KillerRed+ mitochondria (top panels) generate an increase in ROS levels (DCFDA, ROS indicator, bottom panels). Mean fluorescent intensity (MFI) of DCFDA list in the inset. (B) Schematic of optogenetic experiments to generate data in Fig. 3A, D-E. Cells expressing mito-KillerRed are photobleached in a specific ROI containing either cytoplasm only (left) or mito-KillerRed+ mitochondria (right). Following photobleaching, cells are imaged over time to quantify the amount of cell division (Fig. 3A) or ERK activity (Fig. 3D-E). (C) Data from Fig. 3A plotted next to an additional control. Quantification of percent cells divided after no stimulation (left column), photobleaching an ROI containing cytoplasm (cyto, middle column) or an ROI containing mito-KillerRed+ mitochondria (mito, right column). Each data point is the average within a field of view per condition (N= 13 experiments). Error bars represent SEM and scale bars are 10 µm. Wilcoxon matched-pairs signed rank test, *$p<0.05$.

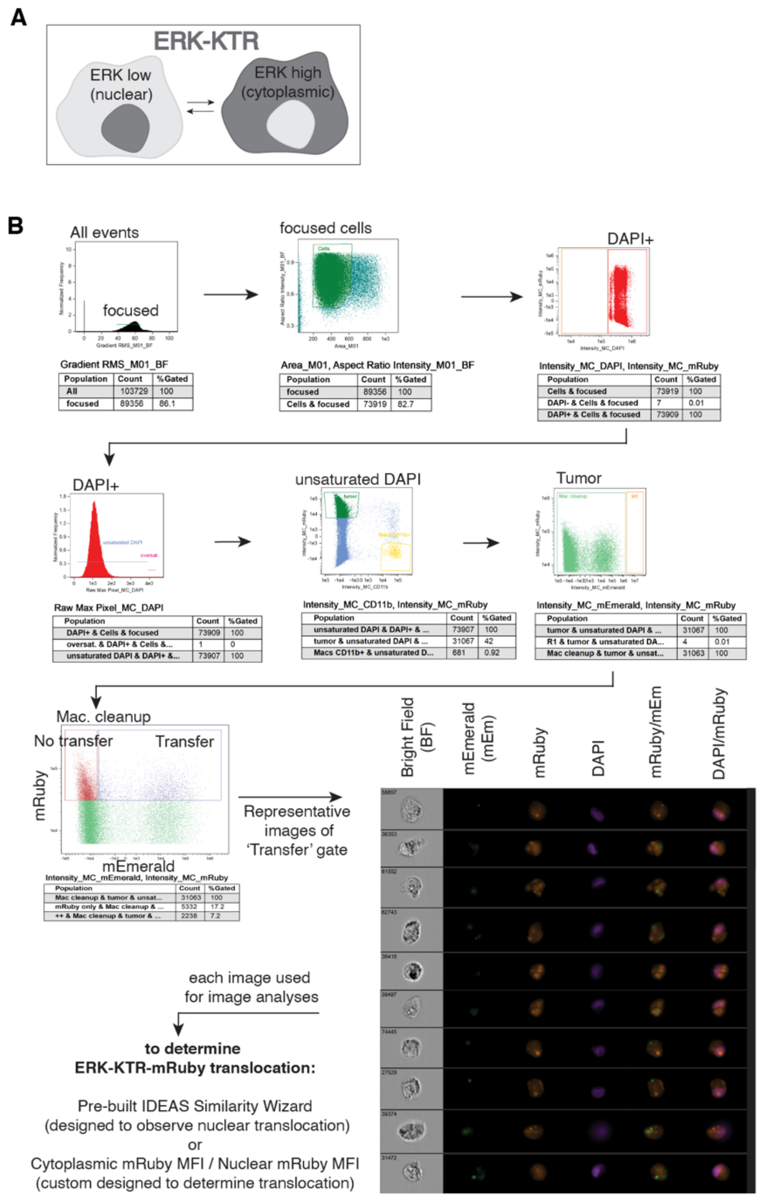


**Fig. S6. Amnis ImageStream pipeline for ERK-KTR quantification.**

(A) Schematic of ERK-Kinase Translocation Reporter (KTR). When ERK activity is low, the fluorescent protein of the KTR resides primarily in the nucleus (left, dark gray). When ERK activity is high, the fluorescent protein translocates to the cytoplasm (right). (B) Workflow of Amnis ImageStream analyses. Mito-mEm macrophages and ERK-KTR-mRuby 231 cells were co-cultured for 24 hours and then analyzed on the ImageStream. Single cells in focus are selected and populations of interest can be isolated for further analyses. Representative images of recipient cells within our "transfer" gate are displayed in all channels acquired (bottom right images). All populations of interest are then put though two independent readouts of KTR translocation: the pre-built IDEAS Similarity Wizard and/or the custom-built cytoplasmic:nuclear (cyto:nuc) mRuby MFI masking algorithm.

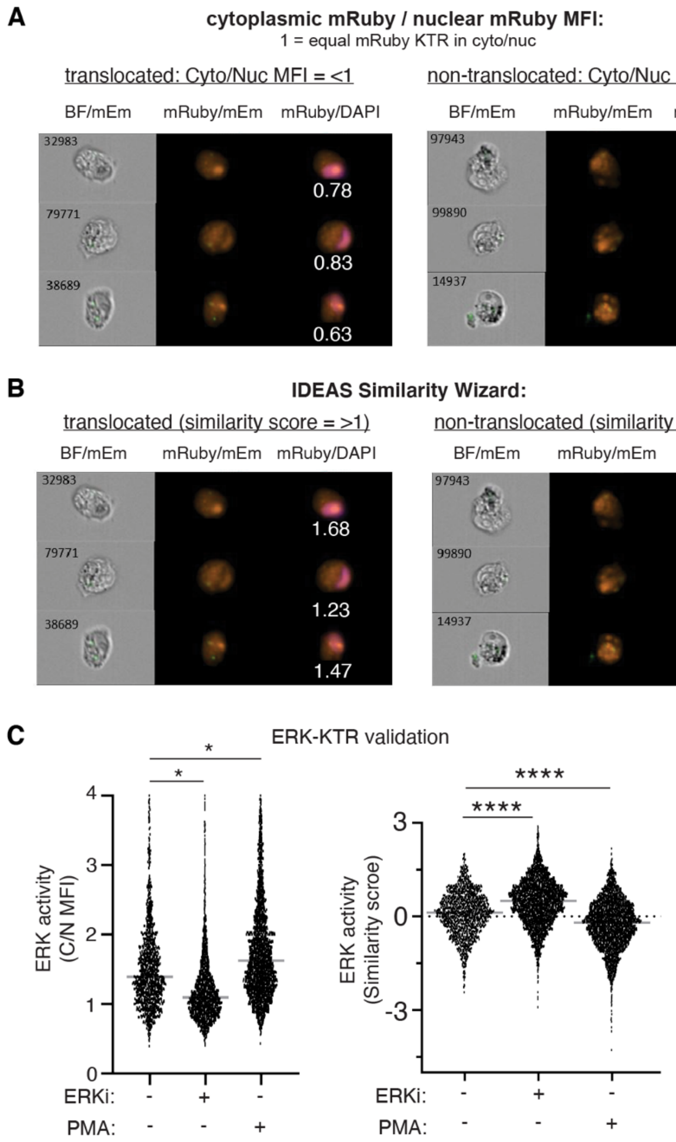


**Fig. S7. ERK-KTR analysis and validation using the Amnis ImageStream pipeline.**

(A) Representative ImageStream images using the mRuby mean fluorescence intensity (MFI) masking algorithm to quantify the amount of cytoplasmic (cyto, C) and nuclear (nuc, N) ERK-KTR-mRuby. Representative images of translocated (left) and non-translocated (right) ERK-KTR images are displayed with the ratiometric cyto/nuc values listed below the corresponding images. (B) Representative ImageStream images using the IDEAS Similarity Wizard to quantify how similar ERK-KTR fluorescent signal is to the nuclear signal (DAPI). Representative images are displayed showing examples of translocated (left) and non-translocated (right) images with the ratiometric values listed below the corresponding images. (C) Quantification of ERK-KTR translocation in 231 monocultures using ImageStream after treatment with 1 μM of an ERK inhibitor (ERKi), SCH772984, or an ERK stimulator, Phorbol 12-myristate 12-acetate (PMA). Cyto/Nuc MFI of ERK-KTR (left) and similarity scores (right) for the same data set are displayed (N=1 donor). Ordinary one-way ANOVA, *$p<0.05$; ****$p<0.0001$.

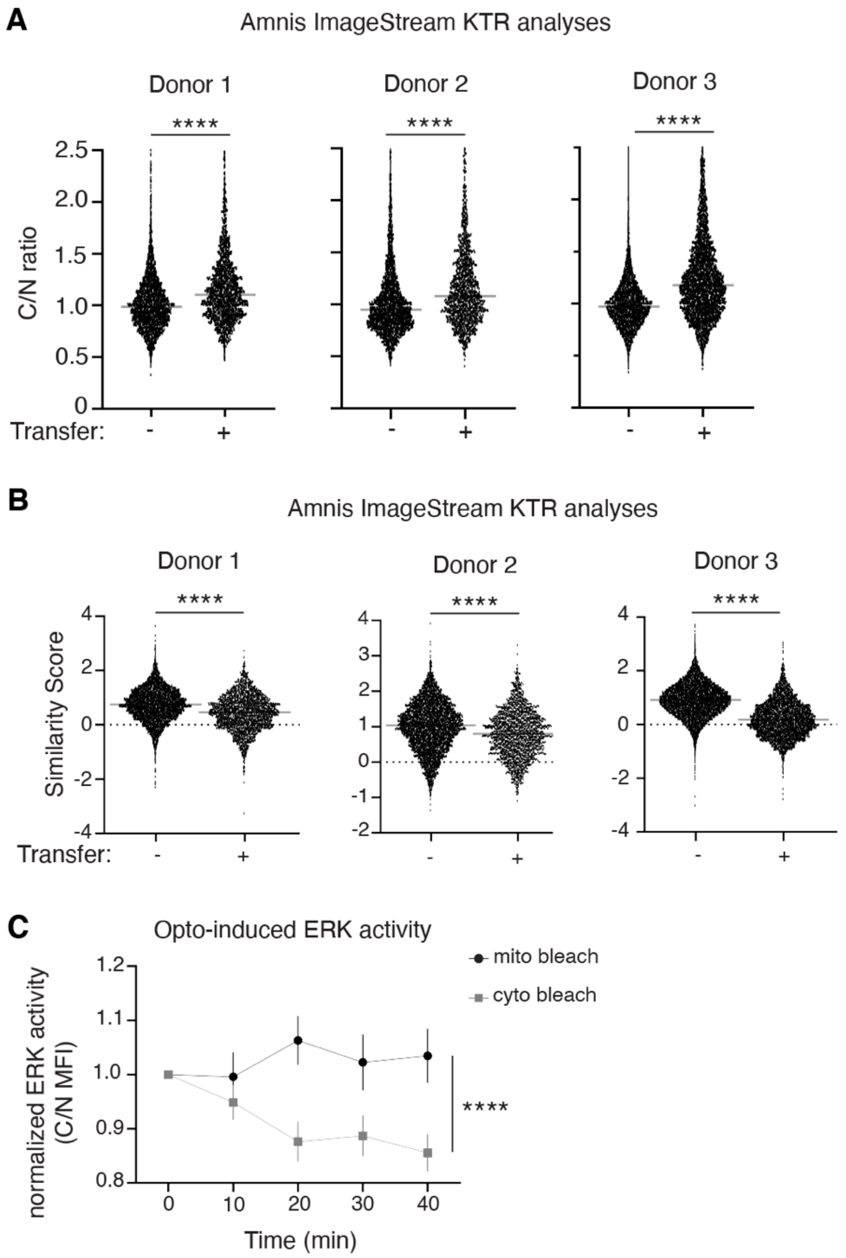


**Fig. S8. Quantification of ERK activity in recipient 231 cells or upon ROS induction**
(A) Cyto/Nuc mean fluorescence intensity (C/N MFI) ratios of the ERK-KTR in co-cultured 231 cells are displayed for 3 different macrophage donors (N=3 experiments). Each data set shows an increase in C/N MFI ratios in recipient cells (left column on each plot) indicating higher ERK activity compared to their non-recipient co-cultured counterparts. (B) The same 3 experimental samples from the analyses in (A) were analyzed using the IDEAS Similarity Wizard. This analysis indicates cells that receive transfer (left column on each plot) have a lower similarity score, indicating higher ERK activity compared to cells that did not receive transfer (N= 3 donors, 3 experiments). (C) Quantification over time of ERK-KTR translocation after mito bleach (black circles) compared to control cyto bleach (gray squares). Unpaired t-test, (A, B) ****p<0.0001. Two-way ANOVA (C), ****p<0.0001.

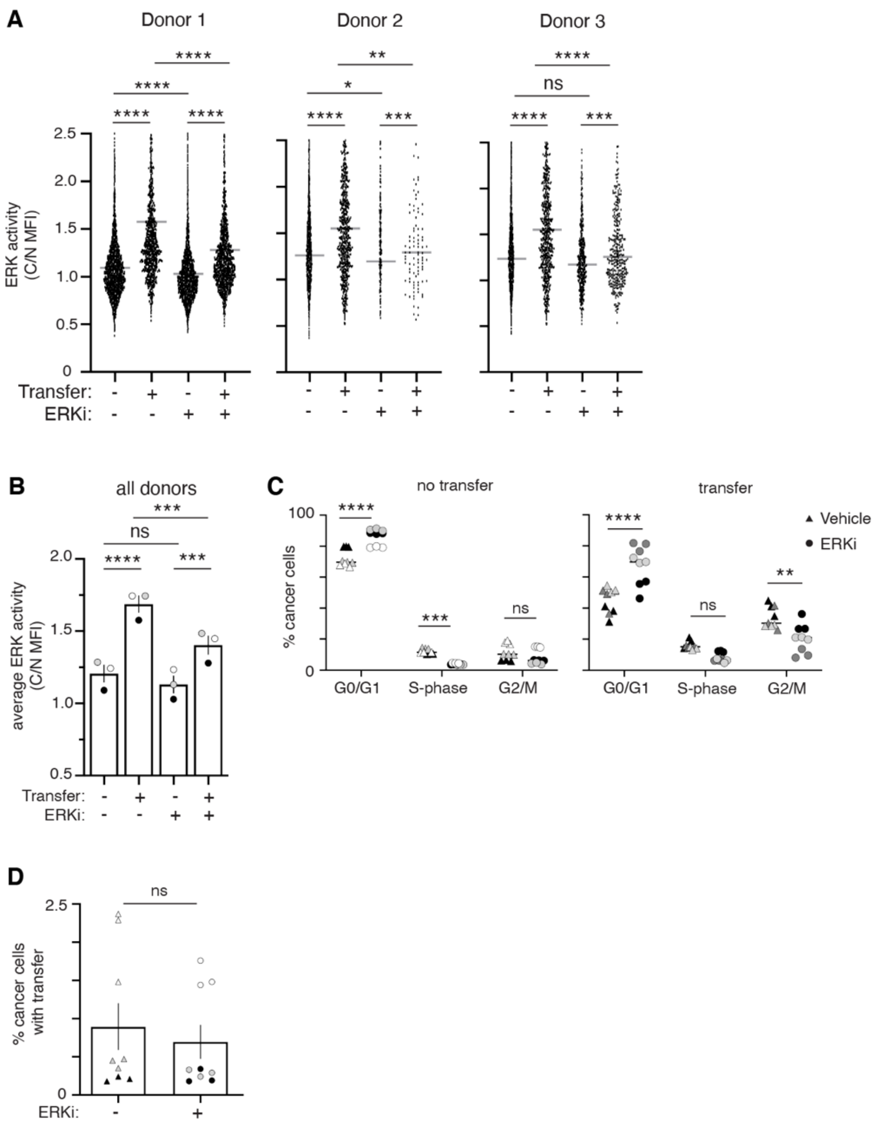


**Fig. S9. ERK inhibition reduces proliferation in cancer cells with macrophage mitochondria**

(A) Co-cultured 231 cells treated with 1µM ERKi or vehicle at time of plating were analyzed for ERK-KTR translocation. Data are displayed from 3 different macrophage donors (N= 3 experiments). Treatment with ERKi significantly reduced the amount of ERK activity in cells that did or did not receive transfer. (B) Average ERK activity from all experiments shown in (A) is displayed (N= 3 donors). (C) Co-cultured macrophages and 231 cells were treated with 1µM ERKi or vehicle at time of plating. Flow cytometry was used to determine how treatment influenced the cell cycle. Plots compare 231 cells that did (right) and did not (left) receive macrophage mitochondria in each stage of the cell cycle treated with vehicle (triangles) or ERKi (circles) (N= 3 donors). (D) Co-cultured 231 cells were treated with ERKi or vehicle at time of plating and the rate of mitochondrial transfer was determined with flow cytometry (N= 3 donors). For all panels, individual donors are indicated as shades of gray and error bars represent SEM. 1-way ANOVA (A), 2-way ANOVA (B-C), Unpaired t-test with Welch's correction (D), *p<0.05; **p<0.01; ***p<0.001; ****p<0.0001.

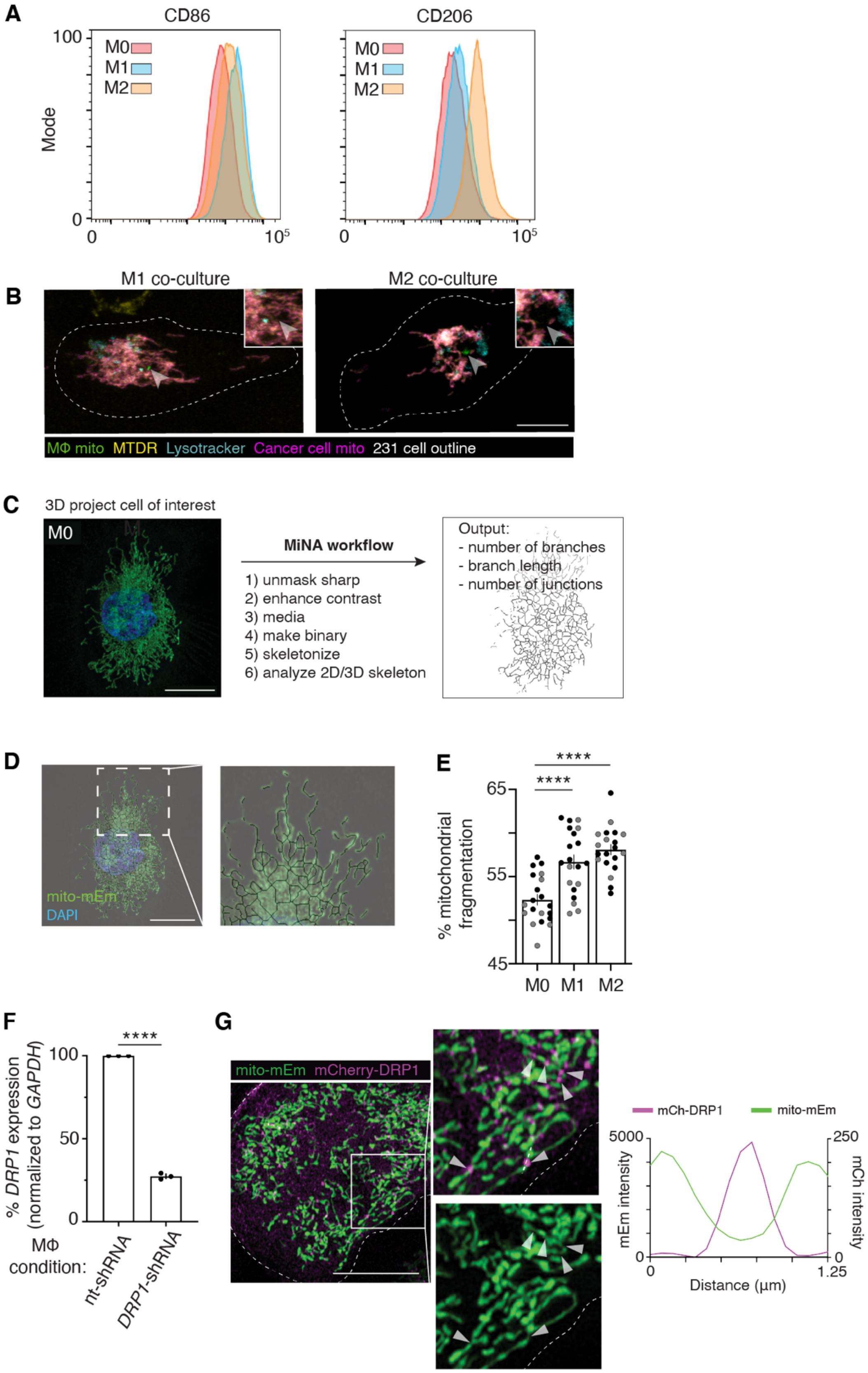
A
CD86
CD206
100
Mode
0
M0
M1
M2
0
$10^5$
B
M1 co-culture
M2 co-culture
MΦ mito MTDR Lysotracker Cancer cell mito 231 cell outline
C
3D project cell of interest
M0
MiNA workflow
1) unmask sharp
2) enhance contrast
3) media
4) make binary
5) skeletonize
6) analyze 2D/3D skeleton
Output:
- number of branches
- branch length
- number of junctions
D
mito-mEm
DAPI
E
% mitochondrial fragmentation
65
55
45
****
****
M0
M1
M2
F
% DRP1 expression (normalized to GAPDH)
100
50
0
****
MΦ condition:
nt-shRNA
DRP1-shRNA
G
mito-mEm mCherry-DRP1
mCh-DRP1
mito-mEm
5000
250
mEm intensity
mCh intensity
0
0
1.25
Distance (µm)

**Fig. S10. Altered macrophage mitochondrial morphology affects mitochondrial transfer to cancer cells.**

(A) Macrophages were activated with IFN-γ (M1 activation; left) or IL-4/IL-13 (M2 activation; right) for 48 hours and flow cytometry was used to determine expression of canonical M1 (CD86, left) and M2 (CD206, right) markers. Representative histograms shown. (B) Representative confocal images of mito-RFP 231 cells co-cultured with mito-mEm M1-like (left) or M2-like (right) macrophages and stained with MitoTracker Deep Red (MTDR, yellow) and LysoTracker (LT, Teal). 100% of transferred mitochondria (green, arrowhead) from M1 and M2 macrophages are MTDR-negative. 63.6% (from M1) and 48% (from M2) of transferred mitochondria do not co-localize with LT (For M1 MTDR and LT staining, N=15 cells, 2 donors; for M2 MTDR staining, N= 24 cells, 2 donors; for M2 LT staining, N= 17 cells, 2 donors). (C) Mitochondrial Network Analyses (MiNA) workflow with a representative input confocal image (left) and example of skeletonized mitochondrial network (right) to quantify number of branches per individual mitochondria, branch length and number of junctions per individual mitochondrion. (D) Representative confocal image with a post-processed skeletonized version of network overlayed. (E) Percent mitochondrial fragmentation of M0, M1, and M2 macrophages (N= 2 donors). (F) q-RT-PCR of DRP1 knockdown (DRP1-KD) macrophages (N= 3 donors). (F) Representative image of mcherry-DRP1 overexpression (magenta, arrowheads) in mito-mEm (green) expressing macrophages. Line scan (right) of the white dotted line in (G) indicates that mcherry-DRP1 protein localizes, as expected, to regions of mitochondria undergoing fission (arrowheads). For all panels, individual donors are indicated as shades of gray, error bars represent SEM and scale bars are 10 μm. 2-way ANOVA (E), Unpaired t-test (F), ****$p<0.0001$.

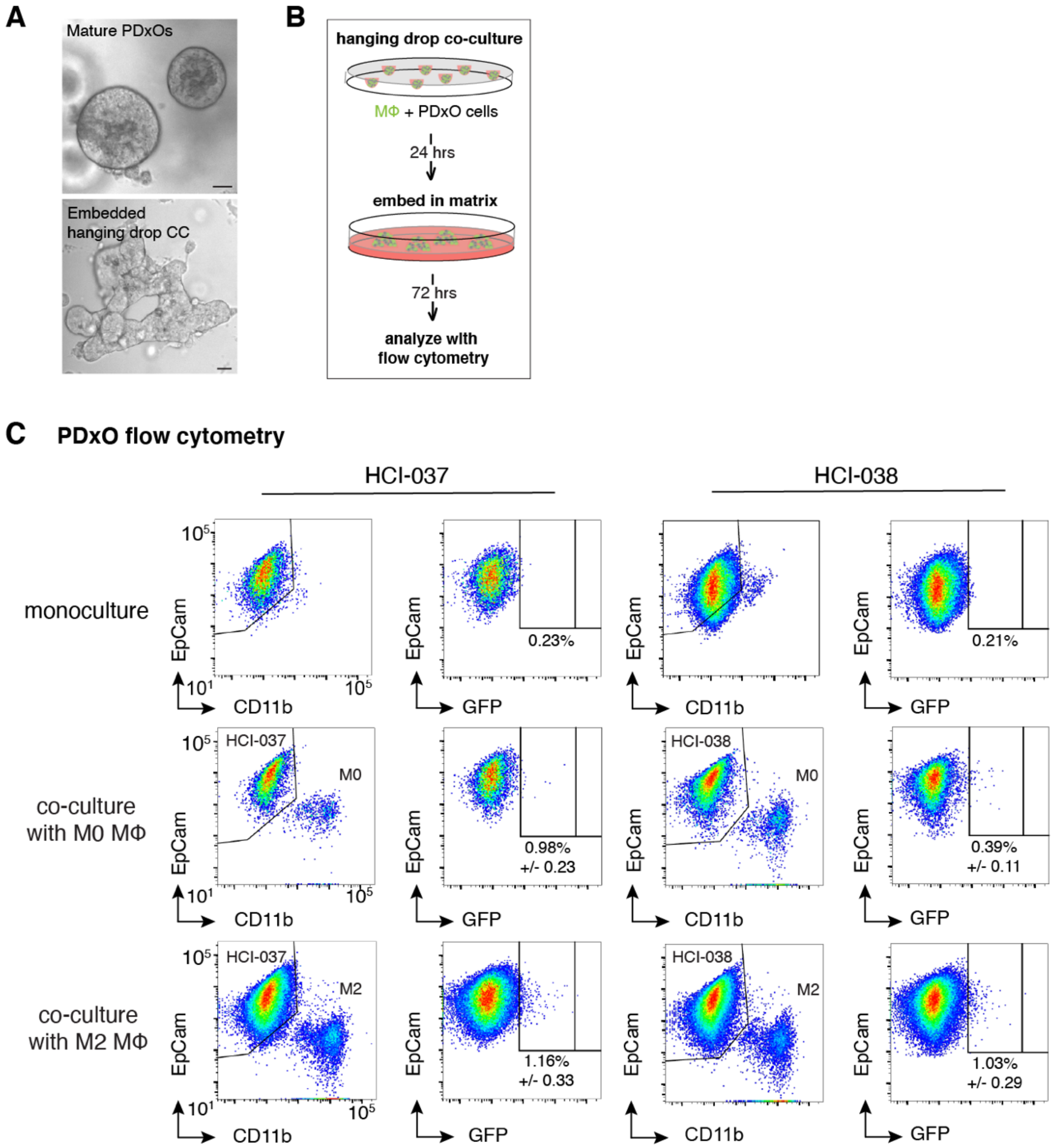


**Fig. S11. Macrophages transfer mitochondria to breast cancer patient-derived cells.**
(A) Representative images of the HCI-037 patient-derived xenograft organoid (PDxO) line in culture (top) or in embedded hanging drop co-culture with macrophages (bottom). (B) Schematic of experimental setup of PDxOs (gray)/mito-mEm macrophages (green) co-cultures. Co-cultures are plated in suspended drops of media (hanging drops) to allow for the formation of cell aggregates without adherence to a substrate. After 24 hours, the co-cultured cells are embedded into a matrix and cultured for 72 hours before analysis with flow cytometry. (C) Top row: Representative flow cytometry plot of PDxO monocultured cells used as the experimental control when quantifying mitochondrial transfer. Middle row: PDxO cells co-cultured with mito-mEM expressing M0 macrophages. Bottom row: PDxO cells co-cultured with mito-mEM expressing M2 macrophages. PDxO lines used for co-culture are indicated at the top of the corresponding panels. Scale bars are 10 µm.

**Movie S1. Macrophage mitochondria are long-lived and remain distinct in recipient cancer cells.**

Movie depicting a recipient mito-RFP expressing 231 cell (magenta) that contains mito-mEm macrophage mitochondria (green in magenta cell, center of frame). 231 cells were co-cultured with macrophages for 7 hours prior to the start of imaging for a duration ~15 hours at a time interval of 5 mins. Maximum intensity projection of images are displayed at 12 frames per second, timestamp in upper left corner in hours (h), and scale bar is 10 μm.
